# Synaptic plasticity facilitates oscillations in a V1 cortical column model with multiple interneuron types

**DOI:** 10.1101/2023.08.27.555009

**Authors:** Giulia Moreni, Licheng Zou, Cyriel M. A. Pennartz, Jorge F. Mejias

## Abstract

Neural rhythms are ubiquitous in cortical recordings, but it is unclear whether they emerge due to the basic structure of cortical microcircuits, or depend on function. Using detailed electrophysiological and anatomical data of mouse V1, we explored this question by building a spiking network model of a cortical column incorporating pyramidal cells, PV, SST and VIP inhibitory interneurons, and dynamics for AMPA, GABA and NMDA receptors. The resulting model matched in vivo cell-type-specific firing rates for spontaneous and stimulus-evoked conditions in mice, although rhythmic activity was absent. Upon introduction of long-term synaptic plasticity in the form of an STDP rule, broad-band (15-60 Hz) oscillations emerged, with feedforward/feedback input streams enhancing/suppressing the oscillatory drive, respectively. These plasticity-triggered rhythms relied on all cell types, and specific experience-dependent connectivity patterns were required to generate oscillations. Our results suggest that neural rhythms are not necessarily intrinsic properties of cortical circuits, but rather they may arise from structural changes elicited by learning-related mechanisms.

## Introduction

Cell-specific cortical activity has been increasingly scrutinized experimentally in recent years, but our mechanistic understanding of cortical dynamics is yet incomplete. Fast (>15 Hz) cortical oscillations are a paradigmatic example, as they constitute a widespread phenomenon but their functional relevance is still under debate (Bosman et al., 2014; Bastos et al., 2015; Vinck and Bosman, 2016; Papadopoulos et al., 2022). Fast cortical oscillations may emerge in neural circuits due to a wide range of mechanistic origins, including excitatory-inhibitory interactions and delayed recurrent dynamics (Tiesinga and Sejnowski, 2009; Bosman et al., 2014). Oscillations have been found to depend not only on simple excitatory-inhibitory interactions (Wilson and Cowan, 1972; Brunel, 2000; Brunel and Wang, 2003), but also on interactions between different cell types such as parvalbumin-positive (PV), somatostatin-positive (SST) or vasoactive intestinal peptide-positive (VIP) interneurons (Rudy et al., 2011; Tremblay et al., 2016), which modulate their emergence as well as other dynamic properties of circuits (Cardin et al., 2009; Lee et al., 2013; Litwin-Kumar et al., 2016; Garcia del Molino et al., 2017; Wood et al., 2017; Antonoudiou et al., 2020; Onorato et al., 2023; Beerendonk et al., 2024).

The functions of fast neural oscillations are however unclear. Electrophysiological evidence traditionally established fast oscillations as a plausible mechanism for intra- and inter-areal communication (Fries, 2005; Bastos et al., 2015; Vinck et al., 2016), with recent work broadening or challenging the idea (Battaglia and Hansel, 2011; Mejias et al., 2016; Papadopoulos et al., 2022; Dowdall and Vinck, 2023). For example, computational models showed that frequency-dependent inter-areal coherence may be enhanced without rhythmic input to neural circuits (Mejias et al., 2016), which questions the potential importance of neural oscillations for communication (Schneider et al., 2021; Dowdall and Vinck, 2023). Human noninvasive recordings have established a link between alpha/low beta rhythms in occipital cortex and local inhibition (Jensen and Mazaheri, 2010; Spaak et al., 2012), but it is uncertain whether oscillations are the cause or consequence of such inhibition. Due to how common neural oscillations are and the lack of a clear functionality, it is difficult to discern whether oscillations are simply a byproduct of canonical neural circuits or whether they are linked more fundamentally to brain function. Characterizing which conditions trigger the emergence of rhythmic activity is a key step in this process.

Here, we tackle the above question by building a biologically detailed model of a cortical column of mouse primary visual cortex V1 and studying the emergence of rhythmic activity. Our model incorporates the dynamics of AMPA, NMDA, and GABA-A receptors, and is tightly constrained by state-of-the-art cortical connectivity data (Thomson et al., 2002; Binzegger et al., 2004; Potjans and Diesmann, 2014; Billeh et al., 2020). This includes cell densities and laminar-specific connectivity across four different cell types (pyramidal neurons and PV, SST and VIP interneurons (Rudy et al., 2011; Tremblay et al., 2016)) and five laminar modules (layers 1, 2/3, 4, 5 and 6). We first fitted the model parameters to replicate spontaneous and stimulus-evoked firing rates in vivo, extending previous cortical column models (Potjans and Diesmann, 2014; Albada et al., 2015). Rhythmic activity was notoriously absent from model dynamics, even with strong external input. However, upon introducing spike-timing-dependent plasticity (STDP), broad-band (15-60 Hz) neural oscillations emerged. These oscillations first appeared within feedforward input layer 4 upon stimulation, and then propagated to other layers as observed experimentally (van Kerkoerle et al., 2014). We showed that (i) the frequency and power of oscillations can be modulated via feedforward or feedback input to the cortical column, (ii) all interneuron types, but mostly PV, participate in the generation of fast oscillations, and (iii) oscillations are a consequence of selective, plasticity-driven pairwise reinforcement of synapses. Our results suggest that neural oscillations might not simply be a byproduct of realistic connectivity patterns in canonical cortical columns, and that their appearance may reflects underlying circuit reconfigurations due to synaptic plasticity. Thus, the functionality of oscillations is not bounded by having a realistic connectivity of cortical columns per se, but by selective and distributed experience-dependent plasticity.

## Methods

### Model architecture

The cortical column model (Moreni et al., 2024), shown in Fig. 1, is composed of a total number (N_total_) of 5,000 neurons. This constitutes a small fraction of the size of a real cortical column in V1 (which may be estimated on 80,000 neurons for a cortical surface of 1 mm^2^ (Potjans and Diesmann, 2014)), but it allows us to simulate the model for long periods of time and allow for plasticity rules to gradually changes synaptic weights. The model consists of four cortical layers each containing pyramidal neurons, PV, SST and VIP cells (layers 2/3, 4, 5, 6) and one layer containing only VIP cells (layer 1). Each of the four ‘complete’ layers harboured pyramidal neurons (85% of cells) and inhibitory interneurons (15%) –with the precise proportion of each inhibitory cell type in each layer given by anatomical data (Billeh et al., 2020) and depicted in Fig. 1 as the relative size of the respective inhibitory population (see Supplementary Tables 1 and 2 for more information). All neurons received background noise from the rest of the brain as shown in Fig. 1. The levels of background noise that each type of cell received can be found in Supplementary Table 9, the tuned spontaneous firing rate effectuated by the Poissonian spike generators connected to each group differed amongst cell types.

**Figure 1:**
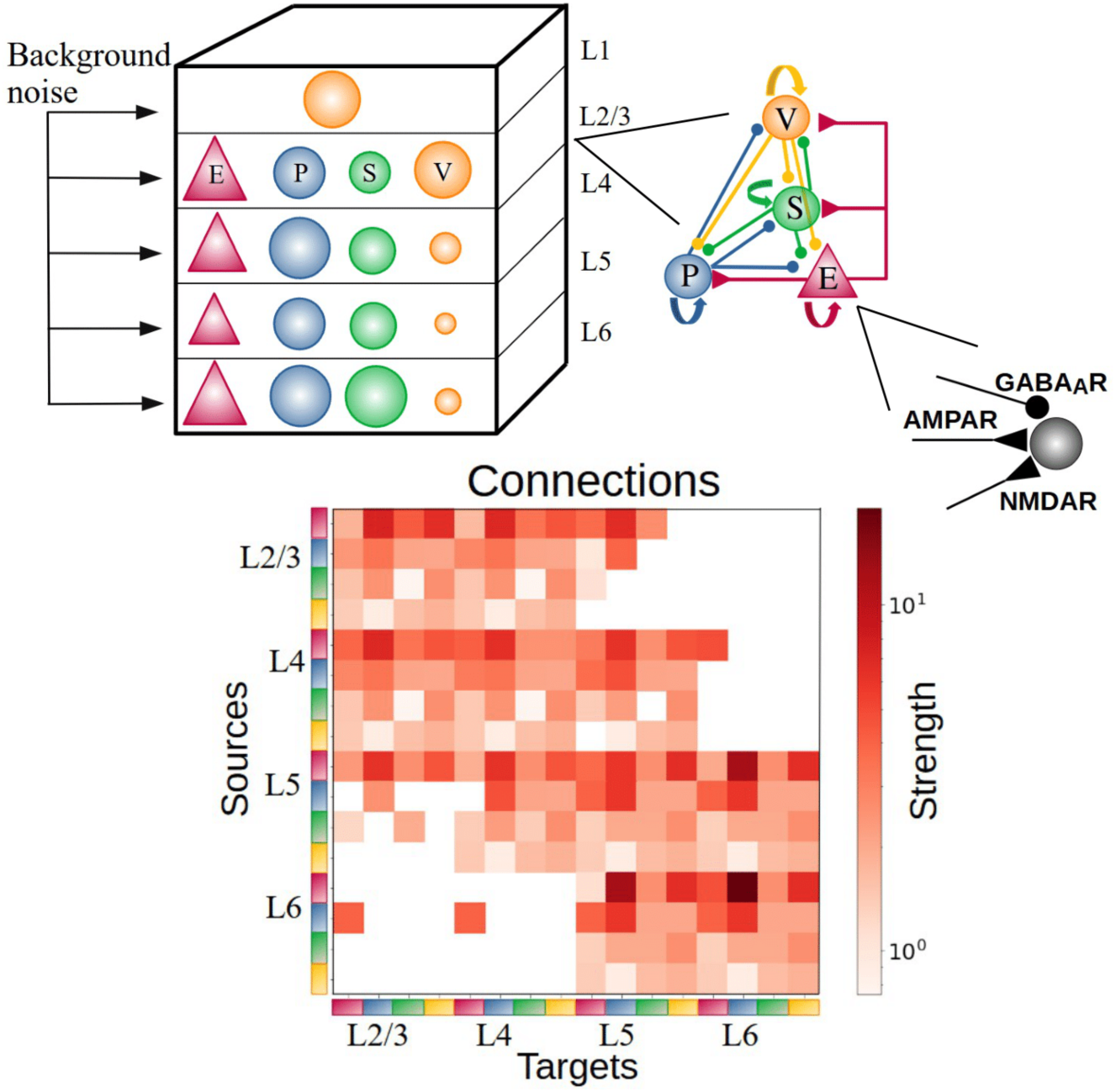
Sketch of the cortical column model. In layers 2/3, 4, 5, and 6 an excitatory population E, (red triangles) and 3 types of inhibitory population (PV, SST, VIP as blue, green, orange circles: P, S and V, respectively) are present. In layer 1 only VIP cells are present. The size of the circles in the top-left panel represents the relative size of the inhibitory populations. Connections between groups are not explicitly shown in the top left diagram; the zoomed-in schematic to the right shows inter-population connectivity and postsynaptic receptors (AMPA, GABA, NMDA) involved. The connectivity matrix is shown at the bottom (adapted from previous work (Billeh et al., 2020)).

### Model for neurons

All pyramidal cells and all three types of interneurons are modelled as leaky integrate-and-fire neurons. Each type of cell is characterized by its own set of parameters: a resting potential *V_rest_*, a firing threshold *V_th_*, a membrane capacitance *C_m_*, a membrane leak conductance g_L_ and a refractory period *1_ref_*. The corresponding membrane time constant is *1_m_= C_m_*/g_L_. The membrane potential *V(t)* of a cell is given by:

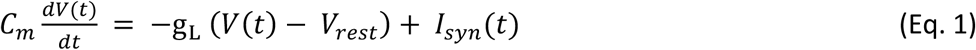

where *I_syn_(t)* represents the total synaptic current flowing in the cell. At each time point of simulation, a neuron integrates the total incoming current *I_syn_(t)* to update its membrane potential *V(t)*. When the threshold *V_th_* is reached a spike is generated, followed by an instantaneous reset of the membrane potential to the resting membrane potential *V_res_*_t_. Then, for a refractory period *1_ref_*, the membrane potential stays at its resting value *V_rest_* and no spikes can be generated. After *1_ref_* has passed, the membrane potential can be updated again (see Tables S5-S9 for the corresponding parameter values).

### Model of synapses

Each cell group in each layer is connected to all the other groups of the cortical column with its own synaptic strength and probability. The values in matrix *P* indicate the probability that a neuron in group A (e.g. a PV cell in layer 4) is connected to a neuron in group B (e.g. an SST cell in layer 5). Excitatory postsynaptic currents (EPSCs) have two components mediated by AMPA and NMDA receptors, respectively. Inhibitory postsynaptic currents (IPSCs) are mediated by GABA_A_ receptors.

The inputs to model neurons consist of three main components: background noise, external (e.g. sensory) input and recurrent input from within the column. EPSCs due to background noise are mediated in the model exclusively by AMPA receptors (*I_ext,AMPA_(t)*) and EPSCs due to external stimuli (i.e. originating from outside the column) are represented by *I_ext_(t)*. The recurrent input from within the column is given by the sum of *I_AMPA_(t), I_NMDA_(t), I_GABA_(t).* These are all the inputs from all the other presynaptic neurons projecting to the neuron under consideration.

The total synaptic current that each neuron receives is given by:

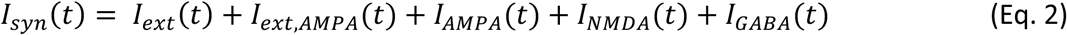

with the last four terms given by

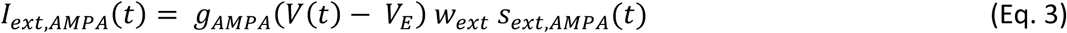

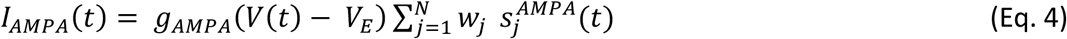

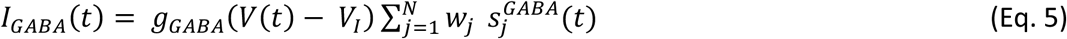

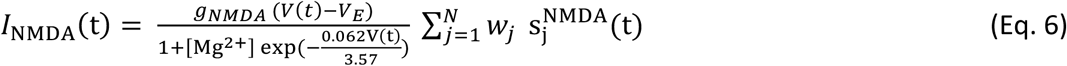

where the reversal potentials are *V_E_*=0 mV, *V_I_ = V_rest_*, and each group of inhibitory interneurons has its own *V_rest_*. The *g* terms represent the conductances of the specific receptor types. The weights *w_j_* represent the strength of each synapse received by the neuron, where synapses transmitting external input to the neurons in the column have a fixed synaptic strength of *w*_)-%_ = 1 (see also Results section). The sum runs over all presynaptic neurons *j* projecting to the neuron under consideration. NMDAR currents have a voltage dependence controlled by extracellular magnesium concentration (Wang, 1999), *[Mg^2+^]=*1 mM. The *s* terms represent the gating variables, or fraction of open channels and their behaviour is governed by the following equations.

First, the AMPAR channels are described by

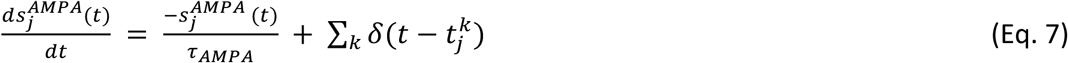

where the time constant of the AMPA currents is 1_AMPA_= 2 ms, and the sum over *k* represents the contribution of all spikes (indicated by delta, *δ*) emitted by presynaptic neuron *j.* In the case of external AMPA currents (Eq. 3), the spikes are emitted accordingly to a Poisson process with rate υ_bkgnd_. Each group of cells in each layer is receiving a different Poisson rate of background noise (see Table S10). The gating of single NMDAR channels is described by

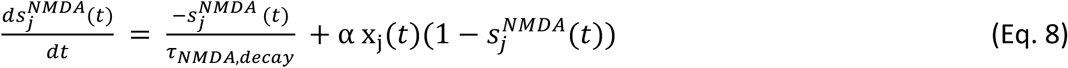

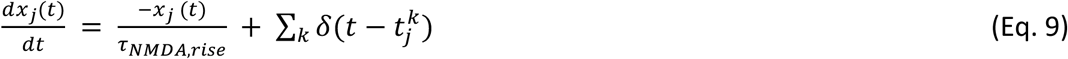

where the decay and rise time constants of NMDAR current are 80 ms and 2 ms respectively, and the constant α=0.5 ms^−1^. The GABA_A_ receptor synaptic variable is described by

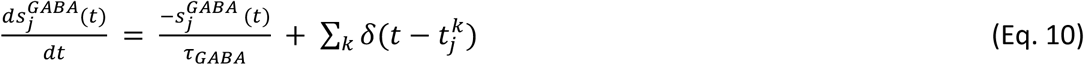

where the time constant of GABA_A_ receptor current is 5 ms.

### Parameters of the model

Each type of cell in each layer is characterized by its own set of parameters: a resting potential *V_rest_,* a firing threshold *V_th_*, a membrane capacitance *C_m_*, a membrane leak conductance g*_L_* and a refractory period *1_ref_*. These data are taken from the Allen institute database (https://portal.brain-map.org/explore/models/mv1-all-layers). In particular, for each type of cell in each layer the Allen database proposes different subsets of cells (e.g. two different PV subsets in L4), each with his own set of parameters. To simplify the model, for each layer we only used one set of parameters for each cell type, choosing the set of parameters of the most prevalent subset of cells. The parameters *C_m,_ g_L,_ 1_ref,_ V_rest,_ V_th_* used for each cell type in each layer are reported in Tables S5-S9.

### External input

In the stimulus-evoked scenarios, specific neuronal populations received external input modelled as a constant current of 30 pA injected at 50% of the neurons in that population. This current represents the simplified visual information transmitted from the retina and the lateral geniculate nucleus. In the particular case of Figure 2, the external input targeted all excitatory neurons in layer 4 instead of 50% of them, to demonstrate the absence of oscillations even in such an extreme case.

**Figure 2:**
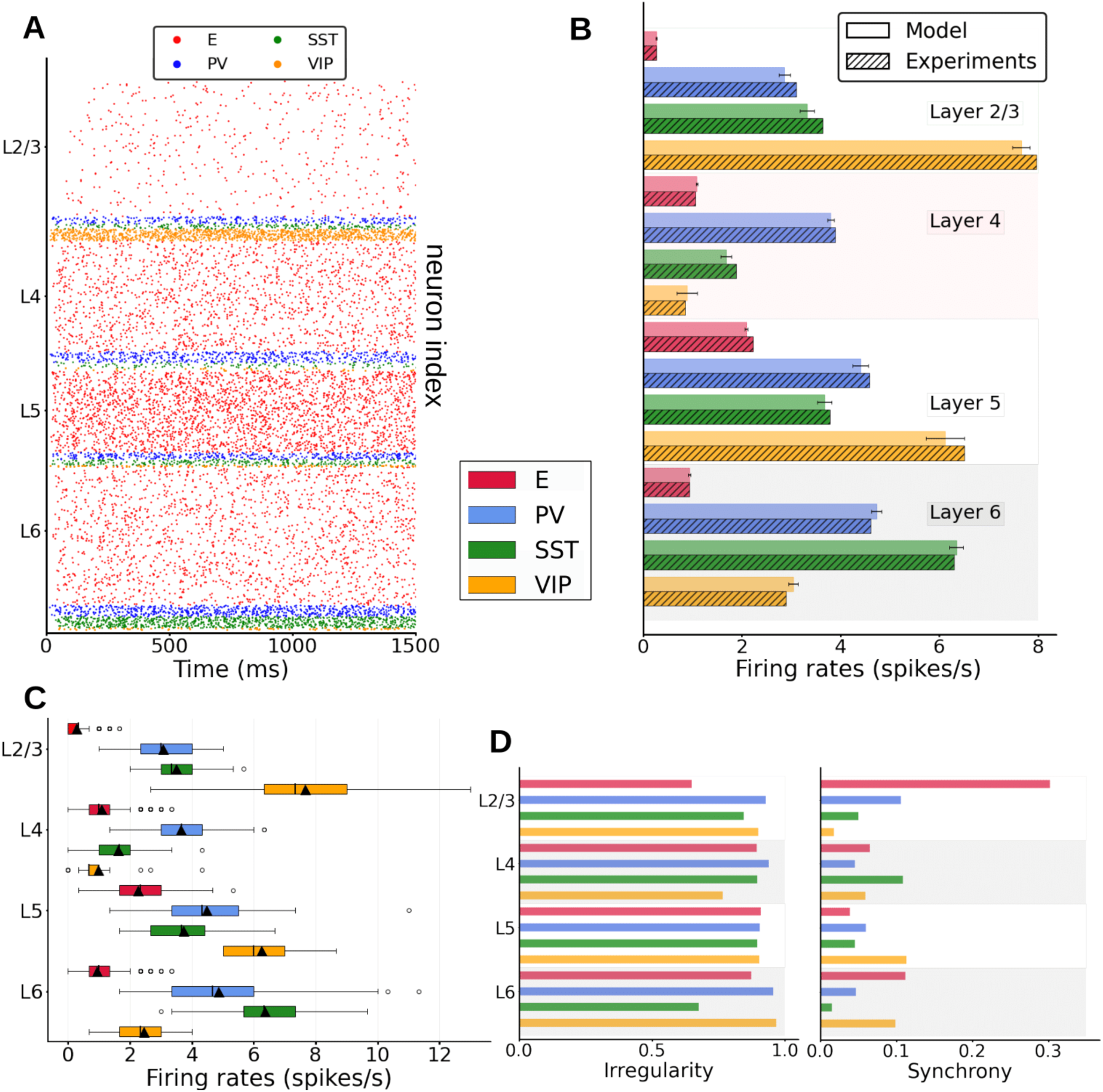
Spontaneous cell-type specific activity in the columnar model. (A) Raster plot of spiking activity simulated for 1500 ms in layers 2/3, 4, 5, and 6 (see inset for cell types). (B) Mean firing rates for each model population (full bars, standard deviation computed over 10 network realizations or initializations) vs experiment (dashed bars, data for SST and VIP cells from Billeh et al., private communication). (C) Boxplot of single-unit firing rates in the model. Circles show outliers, black triangles indicate the mean firing rate of the population, and black vertical lines in each box indicate the median. (D) Left: Irregularity of single-unit spike trains quantified by the coefficient of variation of the inter-spike intervals. Right: Synchrony of multi-neuron spiking activity quantified by membrane potential traces.

### Background input noise

All neurons received background noise, representing the influence of the ‘rest of the brain’ on the modelled area, as shown in Figure 1. Excitatory postsynaptic currents (EPSCs) due to background noise are exclusively mediated in the model by AMPA receptors, denoted as *I_ext,AMPA_(t)* (Eq. 3). The levels of background noise that each group of cells received can be found in Supplementary Table 9. The firing rate υ_bkgnd_ of the background Poissonian pulse generators connected to each group differed among them. Each neuron in every group is connected to its own background Poisson generator. Thus, even though the rates υ_bkgnd_ of the Poisson generators are the same for all neurons within the same group, each specific cell receives its own different pulse train. For the fitting of the external noise values, we (i) manually adjusted the background currents for each isolated population (i.e. G=0, see below), taking advantage of the monotonic input-output relationship for this particular case, (ii) increased the value of global coupling until reaching the desired value, and (iii) readjusted the external background current levels. Steps ii and iii were iterated when necessary. For our specific case, this provided compatible but more refined results than other tested methods like Latin hypercube sampling.

### Synaptic weights

The weight of each synapse w_j_ from neurons of group A to neurons in group B is chosen to be equal to

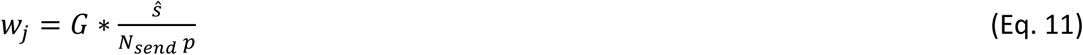

where *G*=5 is the global coupling factor, *s*^ is the overall strength between the two connected groups of cells, *N_send_* is the number of neurons in the sending population A, and *p* the probability of connection between the neurons of the two groups (A and B) taken from the experimental probability matrix *P*. For each pair of connected groups, *s*^ is taken from the experimental synaptic connectivity matrix *S*, defined by the 17×17=289 synaptic strengths between the 17 considered cell types (four groups in each of the full layers, plus one group in layer 1). Matrices *S* and *P* can be found at https://portal.brain-map.org/explore/models/mv1-all-layers. The normalization in Eq. 11 above guarantees that the dynamics and equilibrium points of the system scale properly with the size of the network (Fig. S2).

The spikes generated by an excitatory neuron can target AMPA or/and NMDA receptors of the postsynaptic receiving neuron. The AMPA and NMDA receptors are chosen to be in a 0.8 and 0.2 ratio respectively. Thus, the probability of connection *p* between the neurons of the two groups (A and B) is multiplied by 0.2 for NMDA receptors and 0.8 for AMPA receptors.

### Excitatory synaptic plasticity rule

After having performed simulations with fixed weights, we explored the consequences of allowing synaptic plasticity in the model. To include plasticity we used the STDP learning rule (Gerstner et al., 1996), given by

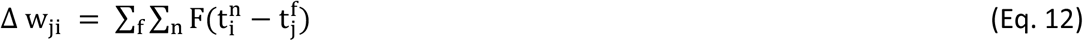

where *n* is the index for the spike times of postsynaptic neuron *j*, and *f* the index for the spike times of presynaptic neuron *j*. The weight change of a synapse depends on the relative timing between pre- and postsynaptic spikes. The function used to account for this change is the following:

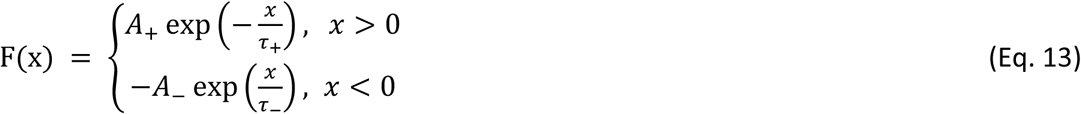

where *x* is the difference between the spike time of the postsynaptic neuron minus the spike time of the presynaptic neuron, and *1****_+_*** and *1****_-_*** are time constants, both with the same value of 20 ms. Likewise, we set the parameters *A****_+_****=*0.02 and *A***_-_**=0.021. The values used to initialize the weights are defined using the *S* matrix (Eq. 11). Only excitatory-to-excitatory weights are allowed to change due to plasticity, and to prevent instability due to ever-growing synaptic weights, we bound them in the range [0, 0.2].

### Inhibitory synaptic plasticity rule

For simulating the network which includes inhibitory plasticity (Fig. S5), the relationship between the modification of the weight *w* of a synapse connecting a pre- and postsynaptic pair of neurons, Δ*w*, and the time difference between their spikes, Δ*t* = |*t*_Q_ − *t*_7_|, is modeled as the difference between two Gaussian kernels:

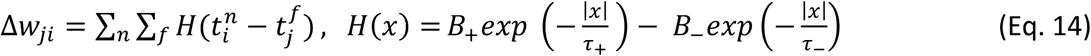

Here i, j are the index of pre and postsynaptic neurons. *τ*_B_(*τ*_A_) is the time constant of synaptic potentiation (depression), taken as 10 (20) ms. *B*_B_(*B*_A_) controls the magnitude of potentiation (depression), taken as 0.04 (0.02).

## Results

### Spontaneous cell type and layer-specific activity

We first match the spontaneous firing rates of all cell types in the model to values observed in vivo. For this, and as done by previous work (Potjans and Diesmann, 2014; Albada et al., 2015), we adjusted the cell-specific background inputs to the neurons and the global scaling connectivity factor. To avoid a fitting simply based on external background inputs, we kept recurrent connections strong enough so that their absence would alter the neural dynamics significantly (Fig. S1A, right panel). The spontaneous spiking activity in the cortical column model (raster plot in Fig. 2A) shows asynchronous irregular activity patterns for all cell types, with firing rates close to in vivo observations (Fig. 2B). We observed a high variability across layers and cell types: pyramidal neurons generally displayed lower firing rates (around 2 Hz for layer 5 and below or close to 1 Hz for other layers), as in the experimental data (Billeh et al., 2020). Firing rates of inhibitory neurons exceeded those of excitatory cells across all layers (except for VIP cells in layer 4), indicating a basal pattern of asynchronous firing in the inhibition-dominated regime (Renart et al., 2010).

Besides the layer and cell-type variability in firing rates values, firing rates of the same type also showed important variability (Fig. 2C): in layer 2/3, individual rates varied from 2 spikes per second to less than a spike per second, in agreement with previous results (Potjans and Diesmann, 2014). Single-unit activity was overall quite irregular, with coefficient of variation values above 0.5 (Fig. 2D, left panel) and standard measurements in membrane potential traces (Golomb, 2007) reflecting asynchronous behaviour.

We then analysed the effects of inactivating different interneuron populations to evaluate their role in controlling the firing activity in the model. Indeed, the activity of all neurons drastically rose when inhibitory neurons were shut down (Fig. S1A, left panel). Furthermore, disconnecting all neurons within the column (which leaves neurons being driven only by the background input) leads to substantial firing rate changes of up to 30% (Fig. S1, right panel), which shows that recurrent connectivity within the column has a notable effect on the activity, even in the spontaneous activity condition being simulated here. We also observed a global rate increase (resp. decrease) in most excitatory (resp. inhibitory) populations, aligned again with an inhibition-dominated network regime.

As a strong cross-correlation between excitatory and inhibitory firing rates is usually indicative of excitatory-inhibitory balance suitable for asynchronous conditions, we have also analysed the cross-correlation of firing rates between excitatory and inhibitory groups in the whole column (Fig. S1B). We observed that the excitatory populations are highly (and positively) correlated with inhibitory firing rate, although this did not correspond to a specific inhibitory group but was rather distributed across different cell populations and even layers. This indicates that excitation-inhibition balance states might be more complex in networks with multiple cell types.

We also performed a finite-size analysis of our model, as some network properties might be hard to estimate from smaller systems (Albada et al., 2015). Our analysis revealed that the firing rates observed in our 5,000-neuron network were close to those obtained with larger networks (up to 20,000 in our simulations, see Fig. S2). This indicates that our model is likely to deliver plausible results for larger and more realistic columnar network sizes, although fundamental limits in reducibility must also be considered (Albada et al., 2015).

### Stimulus-evoked responses of the column

Once the model parameters were fitted to reproduce spontaneous activity, we tested its response to feedforward stimuli mimicking a simple sensory signal. Confronted with a feedforward thalamic input arriving to layer 4, which simulates visual information arriving to V1 from the retina and the lateral geniculate nucleus (Kondo and Ohki, 2016), the columnar model responded with a stereotypical rise and propagation of activity through different layers. Fig. 3A shows an example raster plot of the model when a constant input was given to all pyramidal neurons in layer 4. Pyramidal cells in layers 2/3, 4 and 6 considerably increased their relative activity in response to the stimulus, while those in layer 5 showed a weaker response. Fig. 3B shows the mean firing rates as the input arrives to layer 4 pyramidal cells. Both excitatory and inhibitory populations displayed a significant increase in firing rate due to the input. Inhibitory neurons played a substantial role in how the signal was propagated throughout the column. For example, the reason why pyramidal neurons in layer 5 might respond so weakly to layer 4 input is the PV cell activation in layer 5, where this activity increased significantly (compared to most other inhibitory cell types in other layers) and prevented a further increase in pyramidal neuron activity in layer 5.

**Figure 3:**
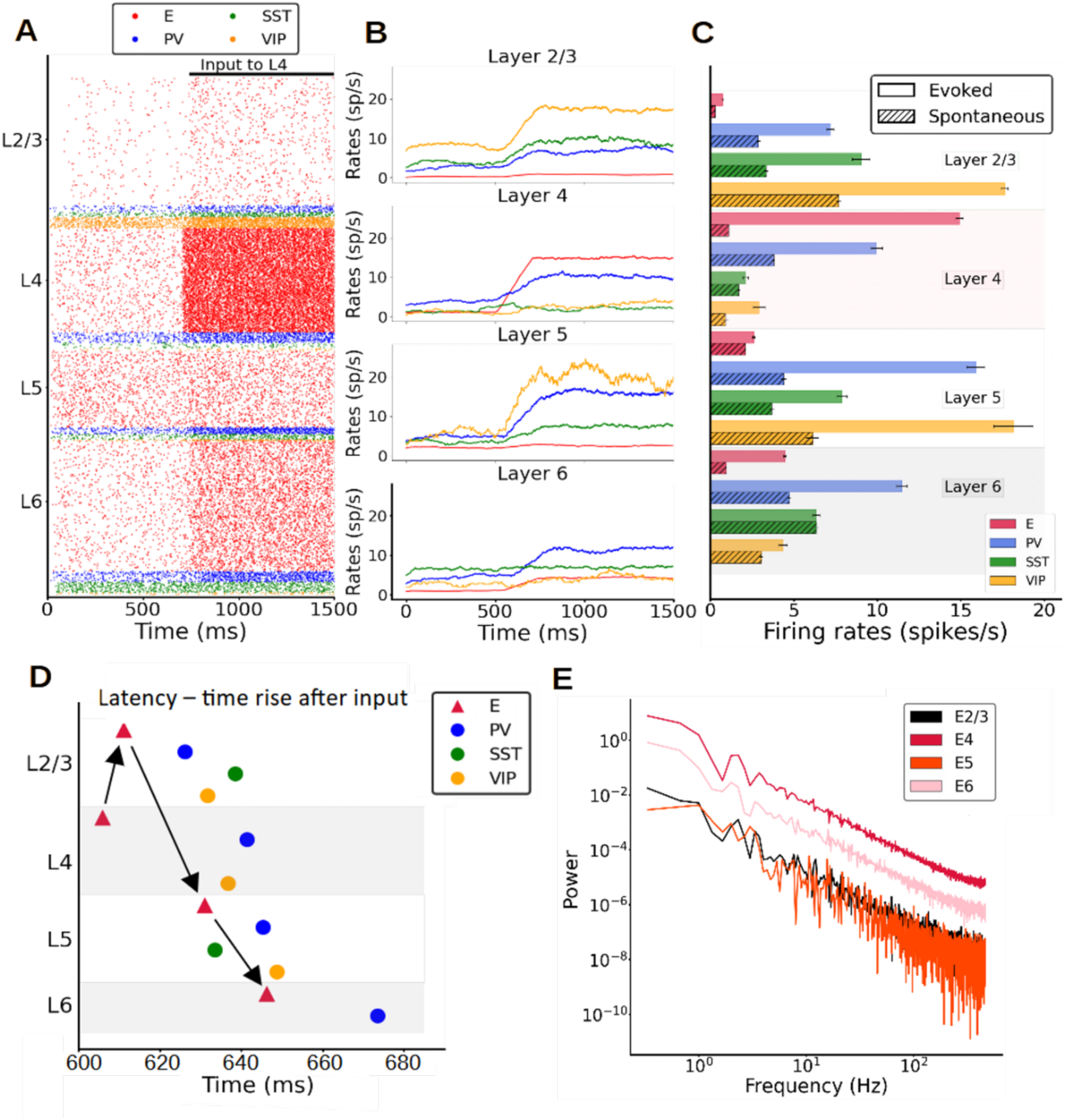
Stimulus-evoked cell-type specific activity in the columnar model. (A) Raster plot of spiking activity for 1500 ms showing the response of neurons after an input current (30 pA) is applied to layer 4 pyramidal neurons at 700 ms. (B) Mean firing rate traces (computed with a 200 ms sliding window and a 1-ms step) showing the increase of the overall activity when the input current is injected. (C) Mean firing rates after stimulus onset of each population for the model (solid bars) vs spontaneous mean firing rates (dashed bars, figure 1B). The error bar for the model are computed as standard deviation over 10 different simulations. (D) Propagation order of the signal elicited by layer 4 excitatory cell stimulation. This is obtained averaging the times rise of 10 different simulations. (E) Power spectrum of excitatory mean firing rates across all layers, showing no signs of oscillatory activity.

To illustrate the power of the model in predicting responses to feedforward stimuli, Fig. 3C shows the mean firing rates after the input had been activated in comparison with experimental data from mouse V1 during passive visual stimulation (Billeh et al., 2020). Although not specifically fitted to reproduce stimulus-evoked activity, the columnar model performed reasonably well and provided good firing rate estimations for all PV interneurons as well as pyramidal neurons (falling within one standard deviation of the experimental range (Fig. S3, (Billeh et al., 2020)). Experimental data on the evoked activity of SST and VIP cells were not available, therefore the results in Fig. 3D may serve as model predictions for future studies.

After the feedforward input excited pyramidal neurons in layer 4, the signal propagated to layer 2/3, then 5 and finally to layer 6 pyramidal cells – generating the sequential activation pattern in canonical microcircuits as proposed in classic neuroanatomical studies (Gilbert, 1983; Douglas and Martin, 2004). This was not captured by previous cortical models (Potjans and Diesmann, 2014; Albada et al., 2015), which indicates that explicitly considering the role SST and VIP neurons is important to understand feedforward activation in the cortical column. The order of responding is clearly shown in Fig. 3D, which shows the activation latency of each population, quantified as the time at which each population reached half of its maximum evoked firing rate. In each layer, the cell type activated first was the pyramidal neuron, followed by inhibitory interneurons after a certain delay, caused by the smoother ramping of inhibitory activity in general. Notably, while the order of activation is the same for pyramidal, PV and VIP cells (layer 4 to 2/3 to 5 to 6), SST cells display a different activation trajectory – increasing their firing rate first on layer 5 and shortly thereafter in layer 2/3, while remaining at spontaneous levels in layer 4 and 6. This suggests that activation patterns in canonical cortical circuits are specific to cell types. The difference in activation latency between the input layer and the deep output layers in our columnar model was about 40 ms, with variations depending on cell type.

While activity in the spontaneous condition was quite irregular (Fig. 2D) as in experimental observations, experimental evidence suggests that rhythmic activity in the range of beta (15-30 Hz) or gamma (30-70 Hz) oscillations is often evoked by visual stimulation. We analysed the temporal evolution of firing rates across all layers in our model and found, however, no evidence of rhythmic activity (Fig. 3E) under a variety of parameter settings, i.e. different external input strengths or sizes of the network. This suggests that the emergence of neural oscillations might require more conditions than explored thus far.

### Introducing synaptic plasticity

After replicating spontaneous activity statistics, our columnar model predicted stimulus-evoked firing rates across cell types and layers, and provided a mechanistic intuition on well-known properties of cortical functioning such as microcircuit communication pathways (Douglas and Martin, 2004) and gain control by deep layers (Olsen et al., 2012). However, so far rhythmic activity was notoriously absent, particularly for stimulus-evoked conditions (Fig. 3E) which would be expected to display stimulus-induced fast oscillations as experimentally observed (Henrie and Shapley, 2005; Ray and Maunsell, 2010; Jia et al., 2013). It is therefore possible that oscillations might not emerge directly from the canonical network of a ‘naive’ cortical column but rather reflect more detailed, experience-dependent changes.

To test this hypothesis, we introduced synaptic plasticity in all excitatory weights of the cortical column model, via the spike-timing-dependent plasticity (STDP) rule (Fig. 4A). To drive long-lasting changes in columnar connectivity, we applied a constant feedforward input of 30 pA arriving at half of the pyramidal neurons in layer 4. Each synaptic weight was initialized as in the results described above. Incorporating STDP resulted in changes in the spiking activity across the column, which evolved from an asynchronous, irregular regime (Fig. 4B, left plot) to a regular, oscillatory pattern only after the plasticity rule led to changes in synaptic strengths (Fig. 4B, middle and right plots). These fast, low-gamma (∼26 Hz) oscillations progressively emerged after STDP was enabled, due to strong, continuous input to layer 4 pyramidal cells. Rhythmic neural activity was present from then on across all layers, with layer 4 displaying the strongest power in the low-gamma frequency band (Fig. 4C). It engaged all cell types to various degrees, as evidenced by firing rate traces (compare e.g. left and right traces in Fig. 4D, corresponding to moments before and after strong changes in the synaptic weight were induced by the STDP rule.

**Figure 4:**
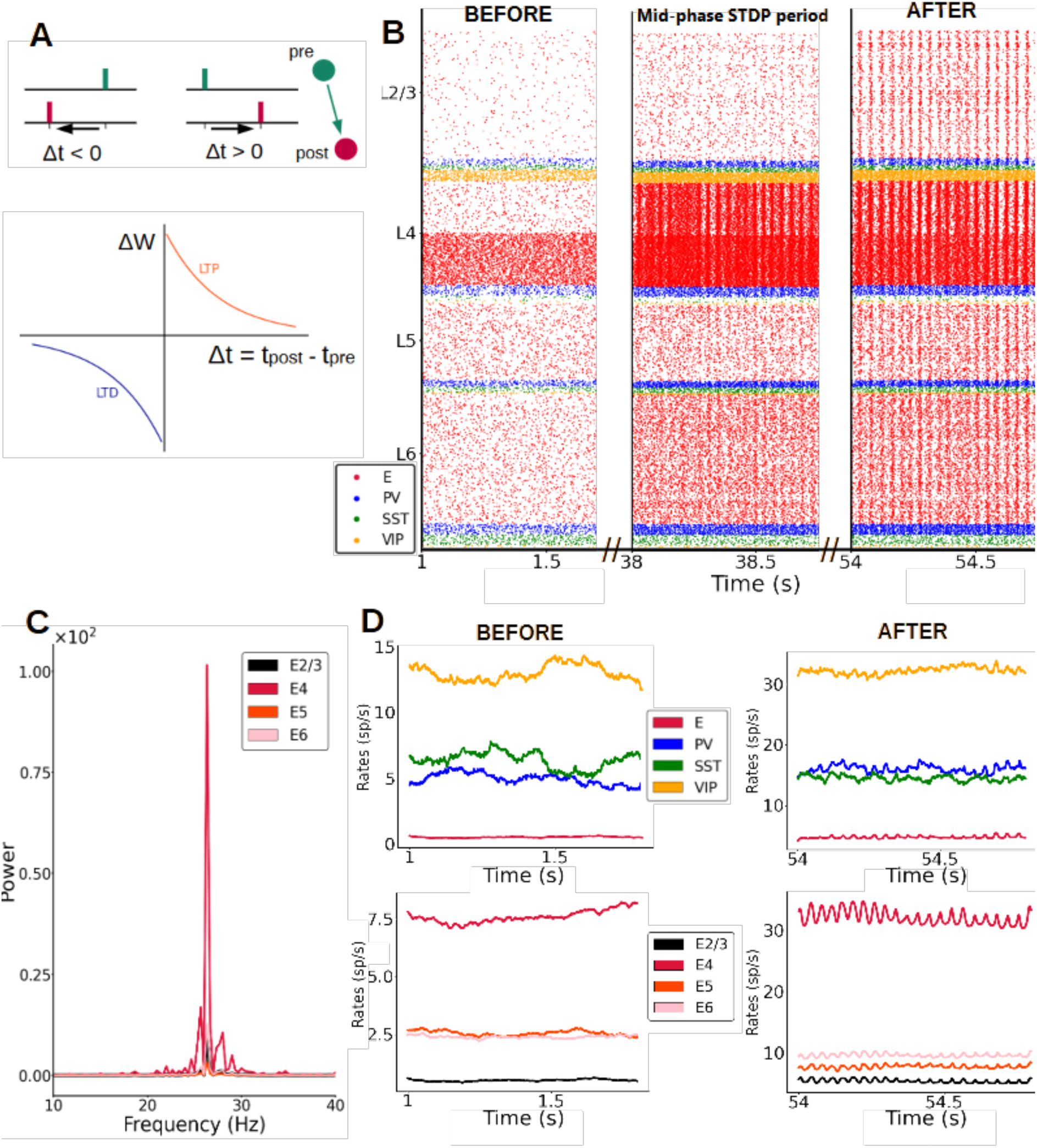
Synaptic plasticity gives rise to fast oscillations. (A) Scheme of STDP rule, with reductions vs increases in excitatory synaptic strength (excitatory-excitatory synapses only) driven by pre- and post-synaptic spike timing. *Δt* is the time between pre and post synaptic spike of a pair of connected neurons. *ΔW* is the change of synaptic weight of the pair according to the STDP rule, LTP stands for long term potentiation and LTD for long term depression (B) Raster plot of spike activity in the column at three example time points after STDP is introduced (concretely 1 s, 38 s and 54 s after plasticity is activated), showing the emergence of oscillations. An input of 30 pA (from t=0.5 s onwards) is given to half of the pyramidal cells in layer 4 and all excitatory-to-excitatory connections in the entire column evolve according to the STDP rule. (C) Power spectrum of the firing rates of pyramidal cells for different layers. The oscillations of pyramidal activity at the end of the plasticity period (55s) have a mean frequency of 26 Hz for an input of 30 pA. (D) Firing rate traces at the beginning (left) and at the end (right) of the plasticity period. Top row: rates in layer 2/3 for all four neuron types. Bottom row: rates of excitatory neurons in all layers.

While most computational studies consider mainly STDP plasticity between excitatory neurons, plasticity in inhibitory synapses has also been observed and modelled and shouldn’t be overlooked (Vogels et al., 2011; Hennequin et al., 2017). We therefore tested our results for a cortical column model in which the network has both (i) the standard STDP plasticity between excitatory neurons, and (ii) an additional rule for inhibitory synapses going from PV cells to excitatory neurons. For the latter, we choose a symmetric “sunken Mexican hat” STDP rule (Luz and Shamir, 2016) (ref “Oscillations via spike-timing dependent plasticity in a feed-forward model”). As Fig. S5 shows, we confirmed our results and observed the emergence of gamma oscillations in this more realistic situation, with the main impact of the inhibitory plasticity rule being a broadening of the power-frequency curve and a decrease of the peak power, especially for the activity in layer 4.

### Oscillatory frequency and amplitude are modulated by feedforward and feedback input

Once plasticity brought synaptic weights into a new stable configuration and the network activity displayed clear oscillations, we fixed the weights to study the resulting dynamics as a function of input strength. We observed that oscillations were triggered and maintained by the feedforward input, and faded away as soon as the input was removed. Importantly, the data-aligned spontaneous firing rates across cell types and layers was conserved in this new modelling scenario (Fig. S4). If the input was switched back on, the oscillations automatically reappeared (Fig. S6). Similar to experimental observations with varying visual contrast (Henrie and Shapley, 2005; Ray and Maunsell, 2010; Jia et al., 2013), the strength of the external input to layer 4 pyramidal cells modulated the frequency and amplitude of the resulting oscillations (Fig. 5A and S7). As the input strength increased, the oscillatory frequency rose from 15 Hz up to 60 Hz (Fig. 5B). The oscillatory power exhibited an inverted-U relationship as a function of the input (Fig. 5C). The effect of (presumably feedback) input to layer 5 had the opposite effect: increasing input strength to layer 5 pyramidal cells (while keeping a constant input of 30 pA to layer 4) led to a reduction in layer 4 pyramidal oscillatory activity, with the reduction being proportional to the strength of input to layer 5 (Fig. 5D and S8). Overall, excitatory input to layer 5 pyramidal cells led to a decrease of oscillatory frequency in all layers except in layer 5, where it increased with input strength (Fig. 5E) and to an overall reduction of oscillatory power across the entire column (Fig. 5F). Thus, feedback input had a strong dampening and slowing effect on the oscillations. Interactions between feedforward and feedback input are therefore able to precisely modulate the rhythmic components of cortical activity.

**Figure 5:**
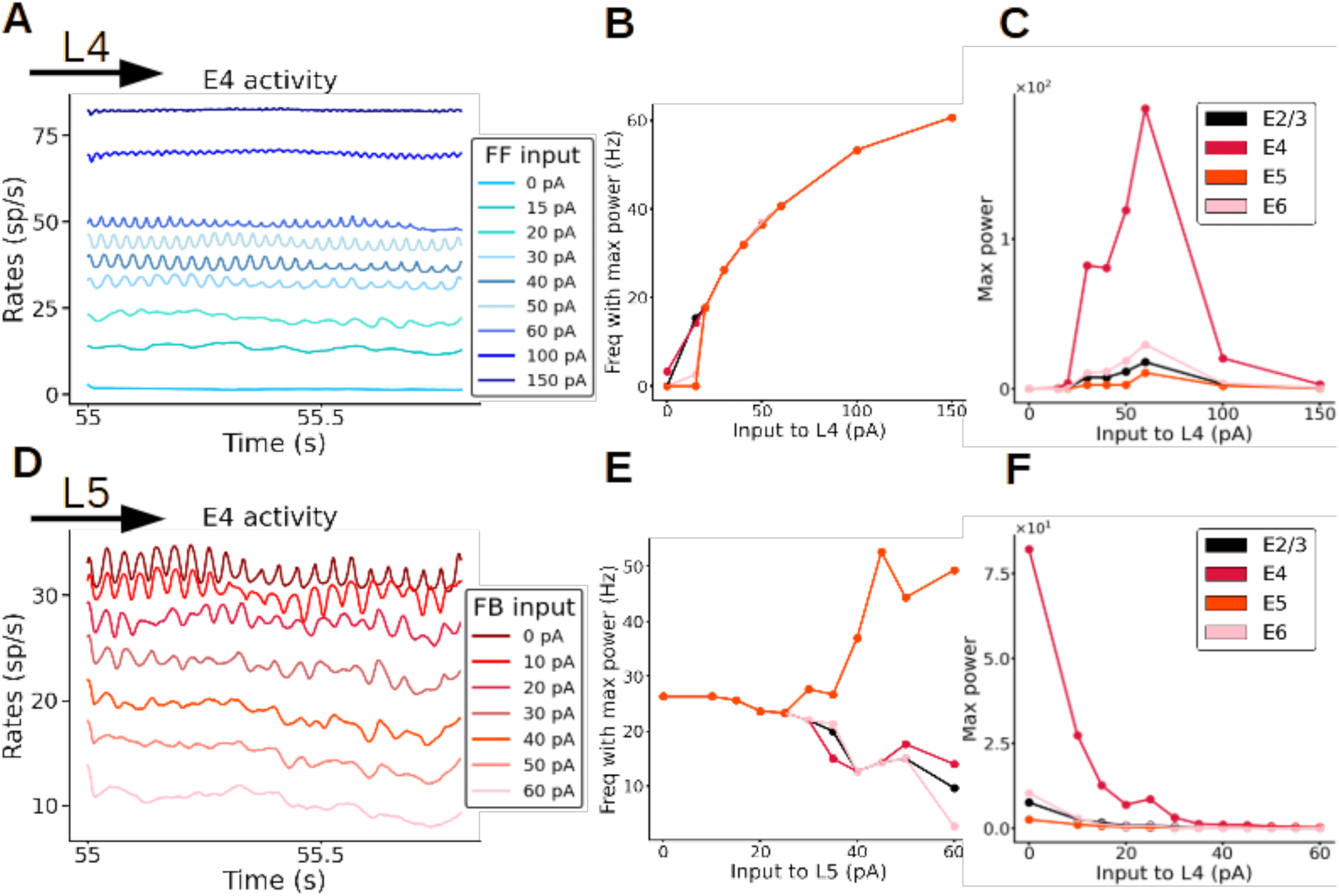
Fast oscillations are modulated by external input. (A-C). Modulation of oscillations of excitatory neurons in layer 4 (E4) by feedforward (FF) input strength. (A) Firing rates of excitatory neurons in layer 4, each colour represents a simulation with a different input strength (colour code in inset). The stronger the input to layer 4, the faster the oscillations are (dark blue trace). When no input to layer 4 is present the oscillations disappear (light blue trace; 0 pA). (B) Frequency of firing rate of excitatory neurons in all layers as a function of input strength to layer 4. (C) Maximal power of oscillations as a function of input strength to layer 4 (D-F) Modulation of oscillations of excitatory neurons in layer 4 by feedback (FB) input strength, while input to layer 4 is kept constant at 30 pA. (D) Firing rates of excitatory neurons in layer 4, each colour represents a simulation with a different input strength to layer 5, (colour code in inset). The stronger the input to layer 5, the slower the oscillations are. When the input strength to layer 5 is very high (60 pA) the oscillations disappear (pink trace). (E) Frequency of oscillatory activity of excitatory neurons as a function of the input to layer 5. (F) Maximal power of oscillations as a function of the input strength to layer 5.

### Origin of cortical oscillations

To better understand the relationship between synaptic plasticity and oscillations as suggested by our cortical column model, it is important to study the mechanisms giving rise to the observed oscillations. Our first aim in this sense was to find out whether oscillations emerge globally and simultaneously across the entire column, or whether a subcircuit directly mediated its generation, driving the rest of the columnar network. A candidate for such a driver role is the layer 4 microcircuit, as it receives the feedforward input initiating experience-dependent processing and has been identified as a gamma generator within visual columns (van Kerkoerle et al., 2014).

When the STDP rule was applied to all excitatory-to-excitatory synapses in the column, the column displayed fast oscillations after a plasticity period as before (Fig. 6A, panel A1). We then repeated the process by allowing STDP-mediated changes in all excitatory-to-excitatory synapses except for those from layer 4, resulting in a network in which oscillations were absent after the plasticity period (Fig. 6A, panel A2). We repeated the process once again, this time only allowing plasticity in excitatory synapses from layer 4. This was sufficient to drive oscillations in all layers (Fig. 6A, panel A3), suggesting that plasticity in the efferent connections from layer 4 is crucial for cortical rhythmicity and that the efferents from layer 4 are the main generator of fast oscillations –which later propagate to other layers, in agreement with experimental data (van Kerkoerle et al., 2014). This also suggests that plasticity in excitatory outputs from layer 2/3, 5 and 6 cells is not crucial for oscillations: without plasticity in these connections, oscillations were still emerging. It is worth noting that the goal of this analysis is to provide more information on the mechanism originating the oscillations, rather than suggesting that layer-specific plasticity would take place in cortical networks.

**Figure 6:**
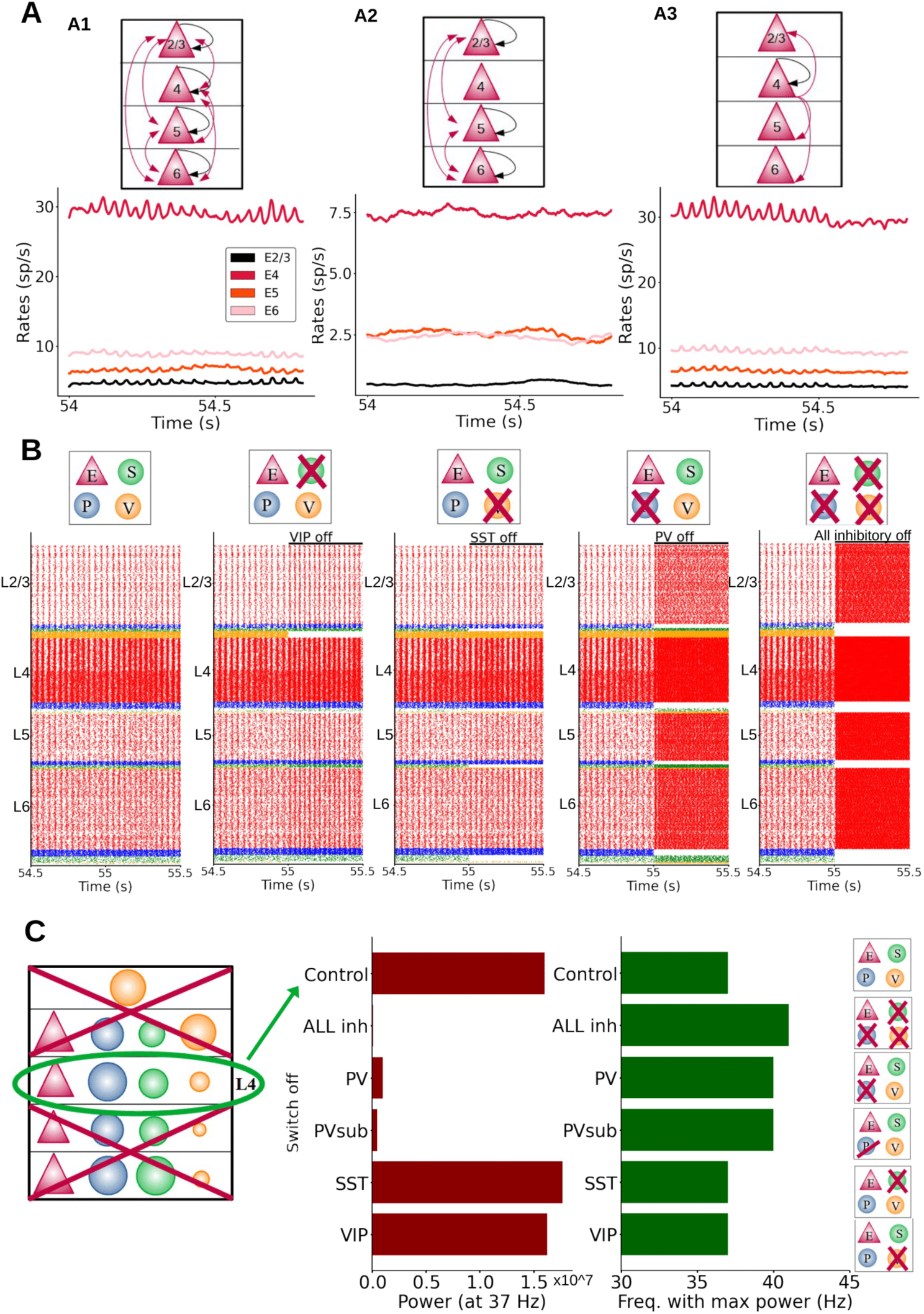
Mechanistic origin of oscillations. (A) Different effects on oscillations depending on the layer in which plasticity of excitatory-to-excitatory synapses is enabled. (Black arrows: plasticity within a group; red arrows: plasticity between different groups). Left: plasticity in all excitatory-to-excitatory connections. Firing rate profiles were obtained while L4 input was provided to half of the pyramidal cells in layer 4. Middle: plasticity in connections between all layers except those from and to layer 4. Right: plasticity enabled only in synapses from layer 4 excitatory neurons. There was no plasticity in the connections going from layer 2/3,5,6 to the other layers. (B) Raster plots of the whole column model for different inactivation conditions. Each group(s) is inactivated at 55 s, to better visualize the effects on neural dynamics. From left to right: control, inactivation of SST, VIP, PV cells and all inhibitory neurons, respectively. Inhibition of PV cells had the largest effect on oscillation frequency and amplitude. (C) Isolated layer 4 circuit. The oscillation frequency in the control situation was 37 Hz. Middle panel: oscillatory power at the 37 Hz frequency for inactivations of different cell groups: all inhibitory, PV, PVsub (same number of PV cells silenced as the number of SST cells present in the column), VIP and SST cells. Note that switching off VIP cells did not change the power of the oscillations drastically as was the case for PV inactivation. Right panel: oscillatory frequency with the maximum power for the same conditions. When switching PV cells off the oscillation frequency (with the maximum power) changed from 37 Hz to a higher value (41 Hz). The power of the oscillation frequency is shown in Fig. S14.

To analyse which cell groups (i.e. a given cell type and layer) are crucial for the emergence of oscillations, we investigated the effect of inactivating different types of interneurons in the whole column. Fig. 6B shows the raster plots for five different conditions, from left to right:

(i) the control condition, and inactivation of (ii) SST cells only, (iii) VIP cells only, (iv) PV cells only, and (v) all inhibitory neurons. Inactivating PV cells had the largest effect amongst specific interneuron inactivations, leading to a sharp increase in the oscillatory frequency across the entire column). Inactivating SST or VIP cells had similar, but much more modest effects on the oscillations. Removing specific groups of interneurons in specific layers led to increases in oscillation frequency (Fig. S9-S13), with the notable exception of layer 5: inactivating interneuron groups in that layer drastically reduced the power of oscillations in other layers, due to the link between layer 5 pyramidal activity and column-wide rhythms (Fig. S10).

Importantly, the more groups we inhibited, the more the oscillation behaviour was influenced. In Fig. S9 we show that removing only one inhibitory group in layer 4 increases the oscillations frequency slightly, with PV having the biggest effect. However, the removal of only one group is not enough to drastically change the frequency of oscillations (Fig. S9). In contrast, silencing more groups at the same time increased the oscillation frequency significantly (Fig. S10). Fig. S11 shows that inhibitory neuron groups in each layer are contributing to oscillations in the whole column, in fact the effect of silencing inhibitory neurons is similar for the different layers (removing inhibition from L2/3 vs L4 vs L6). With these analyses (and more not shown) we can conclude that all inhibitory neuron groups (PV, SST, VIP) make a contribution to maintain oscillations; all of them are relevant even though to different extents (PV being the most important). To further test the relevance of PV interneurons, we silenced all inhibitory groups except one type. Fig. S12 shows that PV cells alone (coupled to pyramidal cells) are able to maintain oscillations, although they became slightly faster. When only SST or VIP cells were maintained, oscillations showed a much higher frequency. This suggests that, while all interneuron types participate in the generation of oscillations, PV cells appear to be more relevant than SST or VIP cells (Perrenoud et al., 2016). Given the important role of PV cells we investigated this further and in Fig. S13 we show how the frequency of oscillations can be modulated by applying differential external input to PV cells in layer 4. The more input current was applied, the slower the oscillations became.

After gaining insight in the full columnar model, we next focused on an isolated subcircuit of layer 4, given its crucial role in rhythm generation. Specifically, we isolated the layer 4 subcircuit by removing all its incoming and outgoing connections with other parts of the column, this was done after the plasticity period had been applied to the entire column from t = 0 until 55 s. This subcircuit was still showing oscillations, although the rhythmicity was not as clear as for the full column. The omission of inhibitory connections from the other layers resulted in faster oscillations: 37 Hz (compared to the full column case: 26 Hz) and a higher firing rate activity. We then inactivated (or silenced) one inhibitory population at a time to observe its effect on oscillatory behaviour. As in the full-column model, we found a particularly strong effect when inactivating layer 4 PV cells, as this severely disrupted oscillations (Fig. 6C and S14). Likewise, inactivating all types of layer 4 interneurons had a strong impact in terms of drops in power, while inactivating SST and VIP cells had only a slight to moderate impact. To test that the impact of PV cells was not due to the higher number of PV cells with respect to SST cells, we also inactivated a subpopulation of PV cells (PV-sub), equal in size to the number of SST neurons in layer 4 (i.e., the largest group of layer 4 inhibitory cells after PV cells), and the results were similar to the inactivation of the full PV population (this was also verified in the full column case, see Fig. S16). The prominent role of PV cells appeared to be related instead of their higher number to the synaptic connections from PV to pyramidal neurons, which are stronger than projections from SST to pyramidal neurons, given the preference of PV cells to establish synapses on pyramidal cell somata (as reflected in the connectivity strength data). Finally, inactivations did not drastically change the layer 4 oscillatory frequency in the case of SST and VIP cells, and led to higher frequencies for PV cells (Fig. 6C and S14). This suggests that, while all interneurons participate in the generation of oscillations, PV cells seem to be more relevant than SST or VIP cells.

### Importance of specific changes in the connectome for the emergence of oscillations

While the above analyses identify some of the key ingredients for the emergence of oscillations in response to long-lasting synaptic changes, it is still unclear what aspects of plasticity are crucial for the emergence of oscillations. It could be that oscillations simply arise due to a global increase (or upscaling) of synaptic weights, or alternatively that refined experience-dependent structural changes due to plasticity play a more important role. To test this, we shuffled the synaptic weights within our cortical model after the plasticity period (Fig. 7A1), as shuffling preserves any overall increase in global coupling level, and observed the impact of shuffling on columnar dynamics. We found that shuffling led to a significant decrease in oscillatory power across all layers (Fig. 7A2 and S17), with oscillations becoming clearly weaker in layer 4 and almost vanishing from other layers such as 2/3 (Fig. 7A3). The oscillatory power for excitatory neurons in layer 2/3 displays a significant drop for its peak frequency (see Fig. S17 for details on all layers). This indicates that specific pairwise reinforcements and the associated structural cross-correlations sculpted by synaptic plasticity are key to the genesis of oscillations.

**Figure 7:**
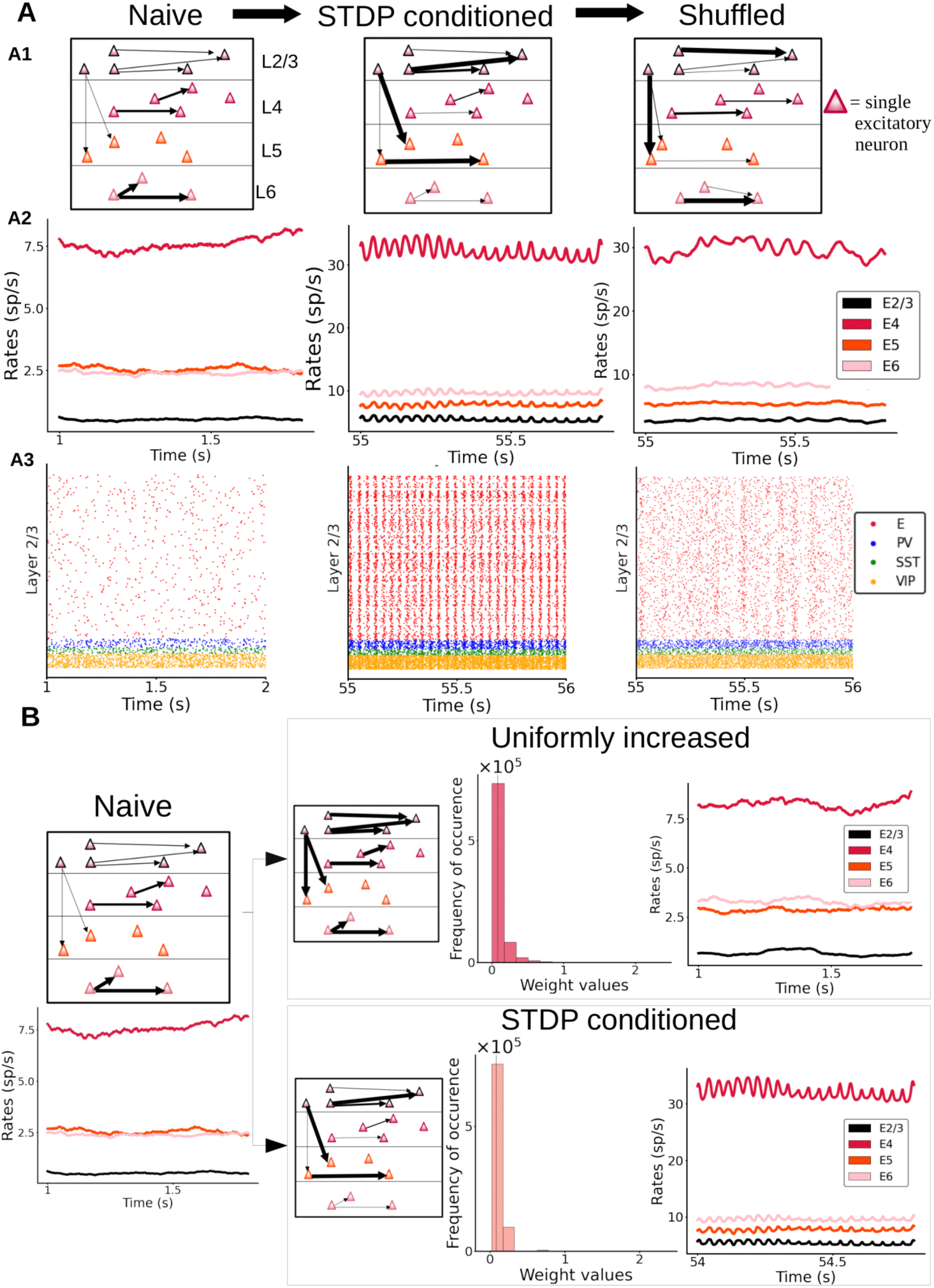
Importance of having a specific distribution of synaptic weights for the genesis of oscillations. (A) Left: “naive” column, prior to plasticity. Middle: columnar model after enabling synaptic plasticity, with activity driven by feedforward input (30 pA to half of the layer 4 pyramidal neurons). Right: same as in the middle panel, but with weights randomly shuffled between individual cell pairs. A2: Firing rate traces of excitatory cells in all layers of the three networks. A3: Raster plots for layer 2/3 are shown for all three networks. (B) A naive column (left) may be either subjected to stimulus-driven changes via STDP (bottom right) or to an equivalent mean increase of all its weights with a similar global weight distribution as the STDP-conditioned network (see histograms with probability distribution of weight values) but without STDP-sculpted structural correlations. Firing rate traces show the presence or absence of oscillations in each case.

Next, we compared the dynamics of a cortical column model with synaptic weights changed due to plasticity (Fig. 7A, middle panels) with a model in which weights were increased, independent from experience-dependent plasticity. In this ‘uniformly increased’ (UI) model, excitatory connections were artificially enhanced from the naive condition to match the overall strength of connectivity of the first model, but without undergoing STDP-regulated changes. This was done by calculating the average synaptic strength of all connections after plasticity induction, comparing this average to the average in the naive network and then increasing the synaptic strength of all connections by that same percentual change in our new UI network. As Fig. 7B shows, a simple increase in global excitatory-to-excitatory synaptic strength is not enough to induce oscillations. This indicates that a specific experience-dependent connectivity pattern and cross-correlations resulting from this pattern are required to generate oscillatory activity in our cortical column model. In other words, the formation of stimulus-driven and experience-dependent ‘resonant assemblies’ is the main driver of oscillations in our cortical column model.

## Discussion

In this work, we built a neurobiologically detailed model of a cortical column of mouse V1, characterizing its dynamics under spontaneous and sensory-evoked conditions and exploring how oscillations emerge in the network. Our model, first and foremost, is built upon detailed empirical datasets of cell-type-specific connectivity patterns between neurons of the visual cortex across all layers. Constraining the model with this data is of the utmost importance to arrive at accurate and relevant predictions. To complete this precise mathematical description of the V1 cortical column, and to more faithfully reproduce rhythmic dynamics and response to input (Brunel and Wang, 2003), we also implemented realistic postsynaptic dynamics for AMPA, GABA and NMDA receptors (Wang, 1999).

For model fitting, we focused on adjusting the parameters for background currents and global coupling strength, as these parameters are difficult to estimate from in vitro recordings but are important determinants of the dynamics (Bernander et al., 1991), and are commonly used for fitting these type of models (Potjans and Diesmann, 2014; Albada et al., 2015). After the parameter adjustments, our columnar model was able to replicate in vivo spontaneous firing patterns (including mean firing rates, spiking irregularity, and synchrony measures) across all cell types and layers. Without further parameter tuning, the model was next able to provide good estimates of activity levels for stimulus-evoked conditions, which are in agreement with existing data for pyramidal and PV cells (Potjans and Diesmann, 2014) and constitute useful predictions for SST and VIP cells. Although other models in the literature have also been able to match firing rates of excitatory and inhibitory cells under spontaneous and evoked conditions, they either did not account for interneuron variability (Potjans and Diesmann, 2014; Albada et al., 2015), readjusted connectivity weights using optimization methods (Billeh et al., 2020), or focused on other brain areas such as barrel cortex (Huang et al., 2022; Jiang et al., 2023). In this sense, our model constitutes an interesting option and an alternative to those models of the mouse V1 column presented until now, particularly for understanding the effects of different types of interneurons on columnar dynamics (Figs. 6 and 7). This also opens the door to use our cortical column model to study the effects of heterogeneity within the same class of cells, using for example existing mean-field approaches (Mejias and Longtin, 2012, 2014).

Our model demonstrated a substantial level of agreement with experimental evidence. Aside from the realistic cell- and layer-specific spontaneous and stimulus-evoked firing statistics, the model successfully replicated a number of experimental observations. First, feeding a feedforward signal to the columnar model triggered the sequential activation of different layers, as predicated by canonical microcircuit diagrams (Gilbert, 1983; Douglas and Martin, 2004). Because previous columnar models without different interneuron types did not show this pattern, this indicates that SST and VIP cells seem to play a role in this translaminar signal propagation – potentially relying on the benefits of interneuronal heterogeneity for signal transmission in cortical circuits (Mejias and Longtin, 2012, 2014). Our model predicts a dependence on SST and VIP cells which can be experimentally tested by optogenetic inactivation. Second, activating pyramidal neurons in layer 5 had an inhibitory effect on other layers, particularly layer 2/3, in agreement with previous observations (Miller et al., 2018). Third, after synaptic plasticity was enabled, feedforward input was able to generate fast oscillations, first in layer 4 and later in other layers. This temporal laminar sequence has in fact been experimentally observed (van Kerkoerle et al., 2014). Fourth, strong feedforward input increased the power and frequency of oscillations (Henrie and Shapley, 2005; Ray and Maunsell, 2010; Jia et al., 2013). Fifth, feedback input suppressed fast oscillations or reduced their power, as observed *in vivo* in V1 recordings (van Kerkoerle et al., 2014).

Besides the generation of the model itself and its validation with existing data, our work hints at a fundamental property of cortical columnar circuits in generating neural oscillations. The precise way by which oscillations emerge in our model is due to emerging neural assemblies. In particular, during strong external stimulation driving the excitatory firing up, the STDP rule allows specific neuron pairs to reinforce their connections, making them easier to synchronize afterwards. Eventually, this neural assembly becomes able to resonate and even spontaneously generate oscillations during stimulation. If the weights are shuffled after the training (Fig. 7A), these detailed neuron-to-neuron correlations are lost, and this leads to a drastic decrease in the oscillations. On the other hand, removing plasticity after the training has taken place (Fig. S6) has no effect on the oscillations, since the correlations have already emerged and are still present. Finally, discarding a leading role of the overall increase of global synaptic weights (Fig. 7B) helps us to conclude that the precise, stimulus-driven association between specific neuron pairs, leading to the emergence of resonant assemblies, is the main driving force behind this phenomenon. These resonant assemblies contain only a small subset of excitatory neurons within the population, which allows to induce oscillatory behaviour without substantially modifying the average firing properties of the whole population. The exact proportion and their relationship with specific interneuron types (in particular, PV cells) should be explored in further work.

The hypothesis that fast oscillations are a direct reflection of experience-driven changes in cortical circuits confronts traditional ideas in the field. Concretely, given that rhythmic activity is ubiquitous in the brain, and that synchrony can be easily obtained in abstract computational neuroscience models, it is sometimes assumed that oscillations naturally arise in canonical neural networks without involvement of experience-dependent plasticity and the fine connectivity structure it entails. Indeed, many models in the literature, some of which are also partially constrained by data, are able to display oscillations without the need of plasticity (Brunel, 2000; Brunel and Wang, 2003; Mejias et al., 2016; Veit et al., 2017; Jiang et al., 2023). However, our work importantly demonstrates that it is possible to reproduce spontaneous and evoked activity with a model that does not display oscillations –which means that rhythmic activity is not necessarily a product of considering realistic canonical microcircuitry. Only by carefully addressing the question with a data-constrained columnar model, as done here, one may reveal the vital role that experience-dependent structural pairwise interactions have on generating cortical rhythmic dynamics. This aligns well with existing ideas which link fast oscillations to flexible mechanisms for the development of cognitive function, such as co-optation mechanisms (Bosman et al., 2014). On the other hand, increasing the strength of all excitatory synapses to high levels in our model would also be expected to lead to oscillations without STDP – synchrony is, after all, an emerging property of networked excitable systems. However, when our model was subjected to STDP, excitatory synapses tended to be weak overall (for example compared to inhibitory neurons), so such a solution, while valid for simple models, does not seem to generalise to more realistic cortical column models.

Despite current neurobiological considerations, our model still presents significant limitations which should be addressed in future studies. For example, the level of biophysical and structural detail for neurons is kept at leaky integrate-and-fire ‘point neuron’ level, although future iterations already in development will expand this to multi-compartment models. Similarly, at the moment our neuron and synapse models lack adaptation mechanisms, such as spike frequency adaptation for neurons or short-term plasticity for synapses. Short-term plasticity would be particularly interesting, given their impact in frequency-dependent communication (Tsodyks and Markram, 1997; Tsodyks et al., 1998; Mejias and Torres, 2009; Mejias et al., 2012) and that fact that recent cortical column models have incorporated such dynamics (Jiang et al., 2023), although specifically for the barrel cortex. Of special importance will be the consideration of long-term plasticity in other synapse types, as we have restricted the majority of the present study to only synapses between excitatory neurons for simplicity, and plasticity rules in excitatory-to-inhibitory or inhibitory-to-excitatory synapses are comparatively less understood. In an effort to provide a first view on the robustness of our results, we have incorporated an inhibitory plasticity rule, a symmetric “sunken Mexican hat” rule (Luz and Shamir, 2016), to our network. Gamma oscillations still emerged in this situation, although with a lower peak power and a broader spectral distribution (Fig. S5). Future studies should be able to incorporate their dynamics using existing descriptions (Vogels et al., 2011; Hennequin et al., 2017), for example to explore the emergence of co-tuning in cortical column models (Lagzi and Fairhall, 2024; Zou et al., 2024). We consider, additionally, that the maximum in synaptic strength imposed in our current model might already approximate, up to some extent, some of the effects expected from plastic inhibitory synapses or homeostatic regulatory mechanisms (Turrigiano, 2008; Vogels et al., 2011), for example limiting the stability problems of excitatory STDP rules (Sjöström et al., 2001; Watt and Desai, 2010).

We have made several experimental predictions based on the results of our model. First, we observed that, while strong positive cross-correlations between excitatory and inhibitory firing rates are found in the model, replicating experimental observations (Renart et al., 2010), this does not always correspond to ‘matching’ pyramidal-PV populations as usually assumed in other models. Instead, we observed a more distributed pattern, where PV, SST and VIP cells within the same or different layers may contribute to match and compensate fluctuations in excitatory firing rates for a given layer. For example, excitatory firing rates in layer 4 are mostly correlated not with local inhibitory populations, but with multiple cell types across other layers. This means that excitatory-inhibitory balance is a far more complex phenomenon in realistic networks with multiple interneuron types, a prediction that could be tested using optogenetics (since electrophysiologically discriminating between inhibitory cell types is challenging). Regarding the activity of different cell types, alternative canonical sequential activations should exist in the V1 column when looking at different interneuron types –for example, upon stimulus onset SST cells will be activated sequentially from deep to superficial layers, while PV and VIP cells will follow the classical sequence just like pyramidal neurons (Fig. 3D). As concerns synaptic plasticity, we predict that plasticity in synapses originating in layer 4 (and targeting other cells inside and outside that layer) will be fundamental for the generation of oscillations in cortical circuits (Figure 6A). Localized pharmacological inactivation of synaptic plasticity mechanisms in layer 4 should reveal important alterations in experience-dependent rhythmic patterns. Although we have not explicitly focused on the role of NMDA receptors in columnar dynamics, the model could be used in future work to study the relationship between pharmacological blocking of NMDARs and alterations in gamma oscillations, or the interaction between NMDAR and different cell types in novel paradigms of working memory (Miller et al., 2018; Mejias and Wang, 2022; Feng et al., 2023). Finally, our inactivation results highlight the crucial role of layer 4 PV interneurons in the emergence of gamma oscillations in the cortical column. Given the relatively strong (weak) projections from PV to pyramidal (SST/VIP) neurons respectively (Fig. 1), and the fact that the same effects are observed in the whole column and in the layer 4 subcircuit (Fig. 6C and Fig. S9), our results indicate that inhibitory synapses from PV to pyramidal cells in layer 4 are particularly important for generating gamma oscillations, a predictions that can also be tested experimentally.

Overall, our biologically realistic, data-constrained cortical column model suggests a clear link between experience-dependent plastic changes and the emergence of neural oscillations. These oscillations are not present anymore if synaptic weights are randomized after learning, and simply upscaling the values of all excitatory synapses in the non-STDP model is also not enough to obtain them. In realistic columnar models, a subtle, self-organizing distributed process driven by experience is needed to produce oscillations.

## Acknowledgments

This project has received funding from the European Union’s Horizon 2020 Framework Programme for Research and Innovation under the Specific Grant Agreement No. 945539 (Human Brain Project SGA3). This work was done with the support of EBRAINS and HBP computing services. We thank Matthias Brucklacher, Kwangjun Lee and Parva Alavian for constructive discussions, and Hans Ekkehard Plesser, Sacha van Albada, Walter Senn and Sander Bohte for their useful feedback on early iterations of the work.

## Author contributions

JFM and CP conceived and designed the study; GM and LZ performed the research; GM, LZ and JFM analyzed the results, GM, LZ, CP and JFM wrote the manuscript.

## Data and materials availability

All information needed to reproduce the results of this manuscript are in the main text and Materials and Methods section, and the code used will be made available upon publication of this work.

## Supplementary figures

**Figure S1:**
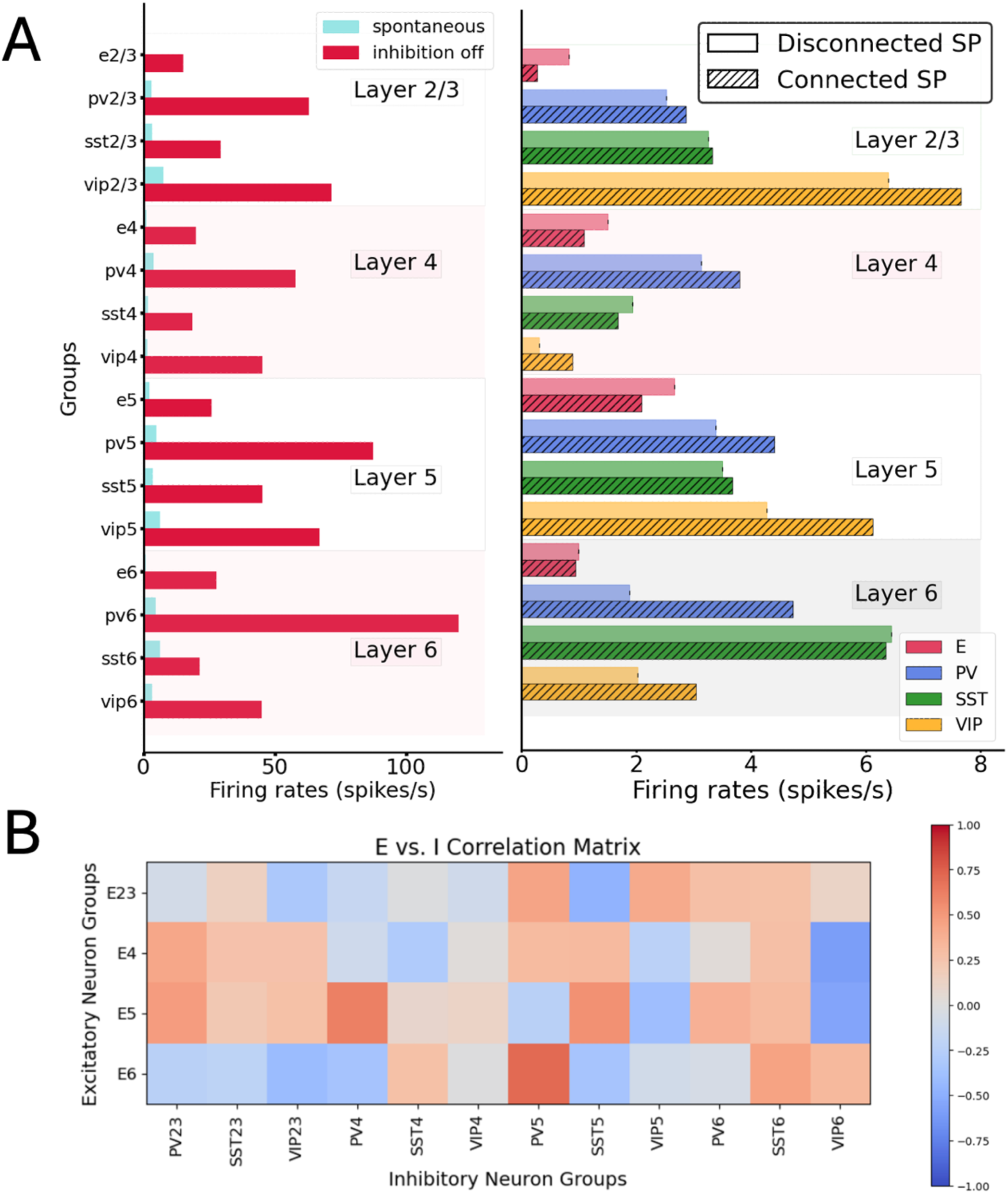
A, left: Comparison between spontaneous activity in normal conditions (control, blue) vs. the condition of disabling the output from all inhibitory neurons (while maintaining their capacity to spike). All groups substantially elevated their firing rates, particularly those in deep layers. A, right: Comparison between spontaneous activity in the condition of connecting and disconnecting all the recurrent connections inside the column. An overall increase (decrease) in all excitatory (inhibitory) neurons were indicative of an inhibition-dominated network behavior. B: cross-correlation matrix between firing rates of excitatory and inhibitory cell groups across the whole column.

**Figure S2:**
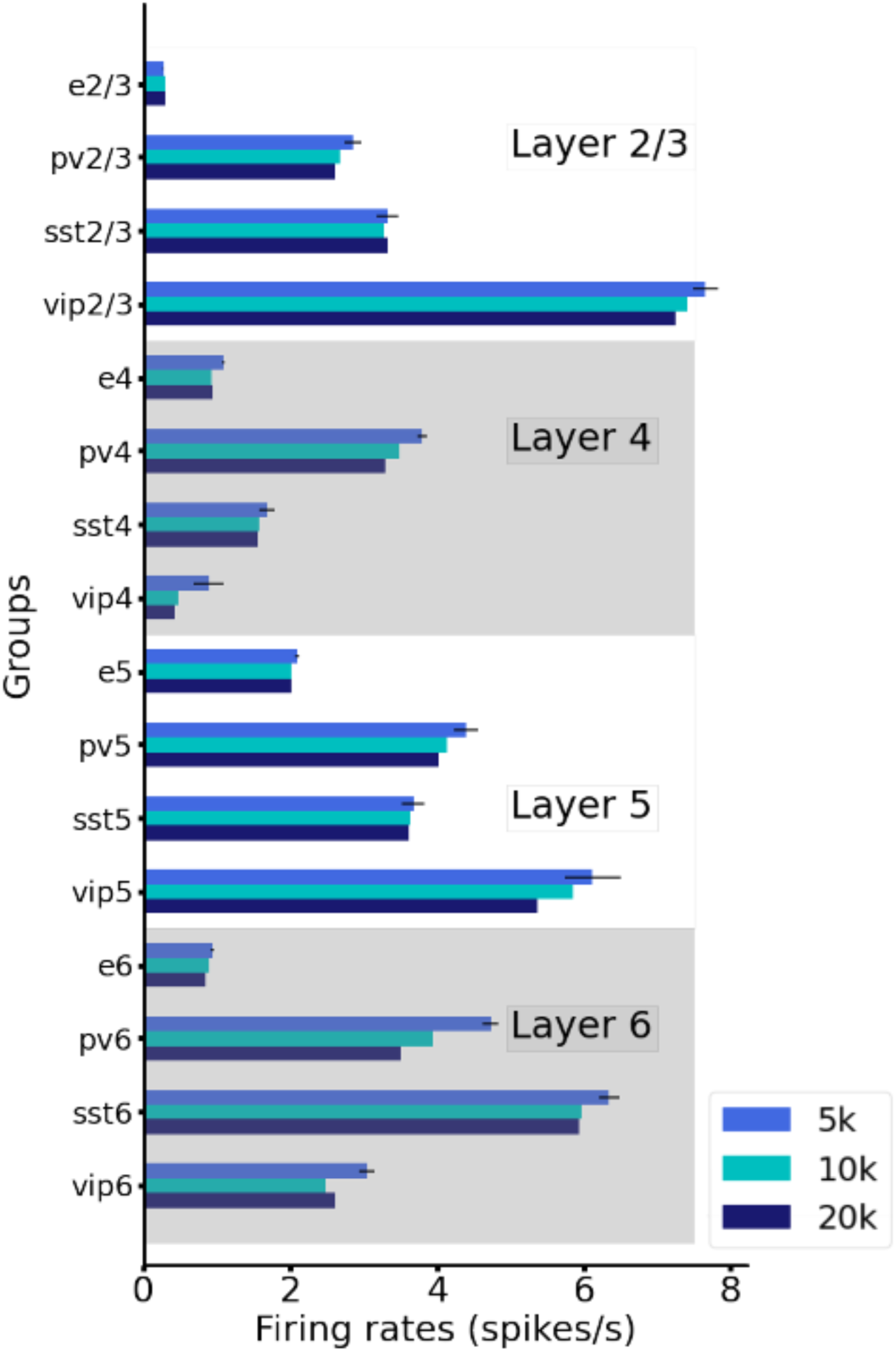
Mean firing rates of all groups for networks of different sizes. We show that, with proper scaling of the weights (see Methods), results for 5000 neurons, 10000 neurons and 20000 neurons lead to the very similar results.

**Figure. S3:**
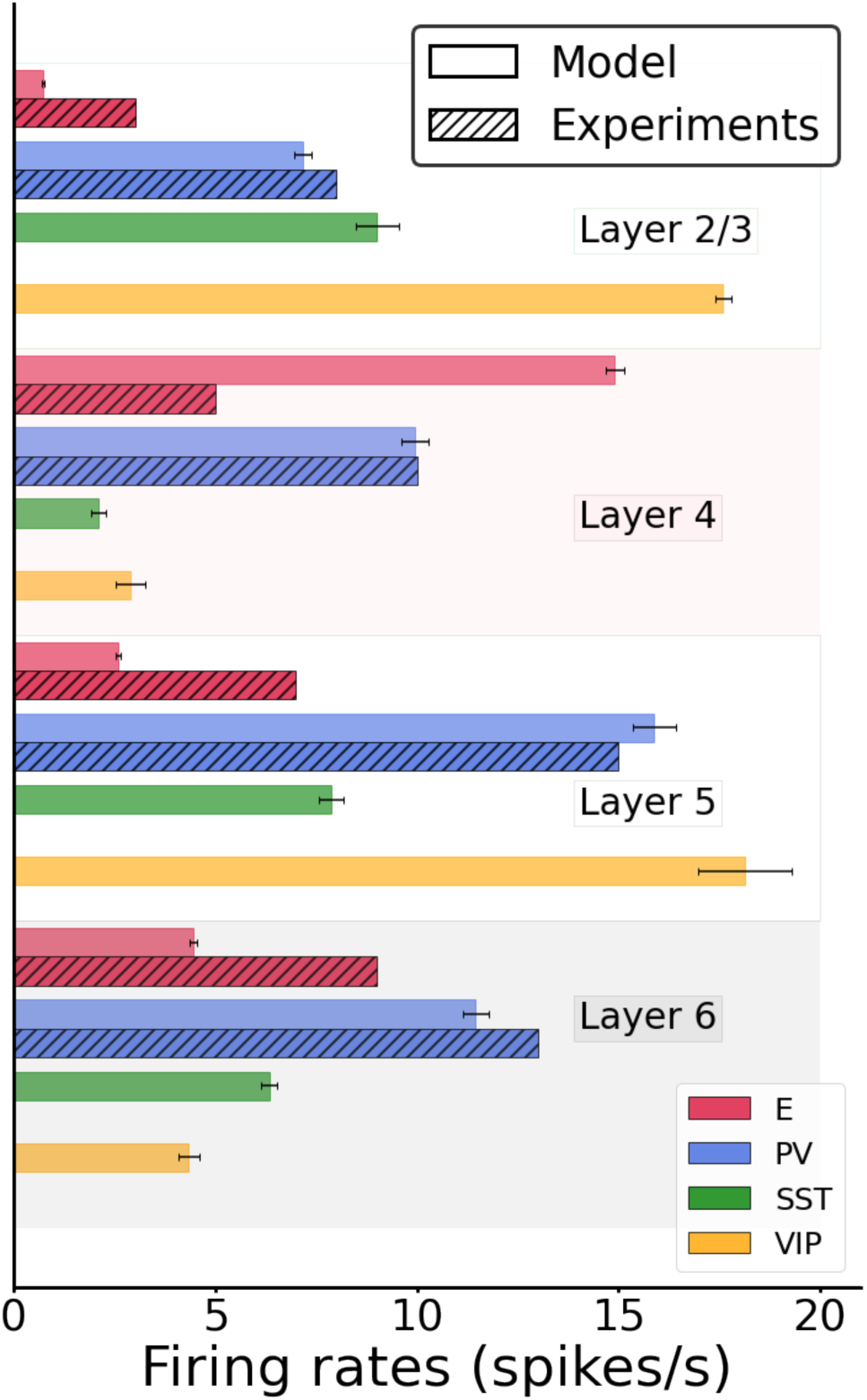
Mean firing rates after stimulating layer 4 excitatory neurons (dashed bars). The error bar for the model are computed as standard deviation over 10 different simulations. Error bars for experimental data are not depicted because of its unrealistically significant variations. Boxplots of experimental firing rates can be found in Billeh et al.^22^, in their figure 3F.

**Figure S4:**
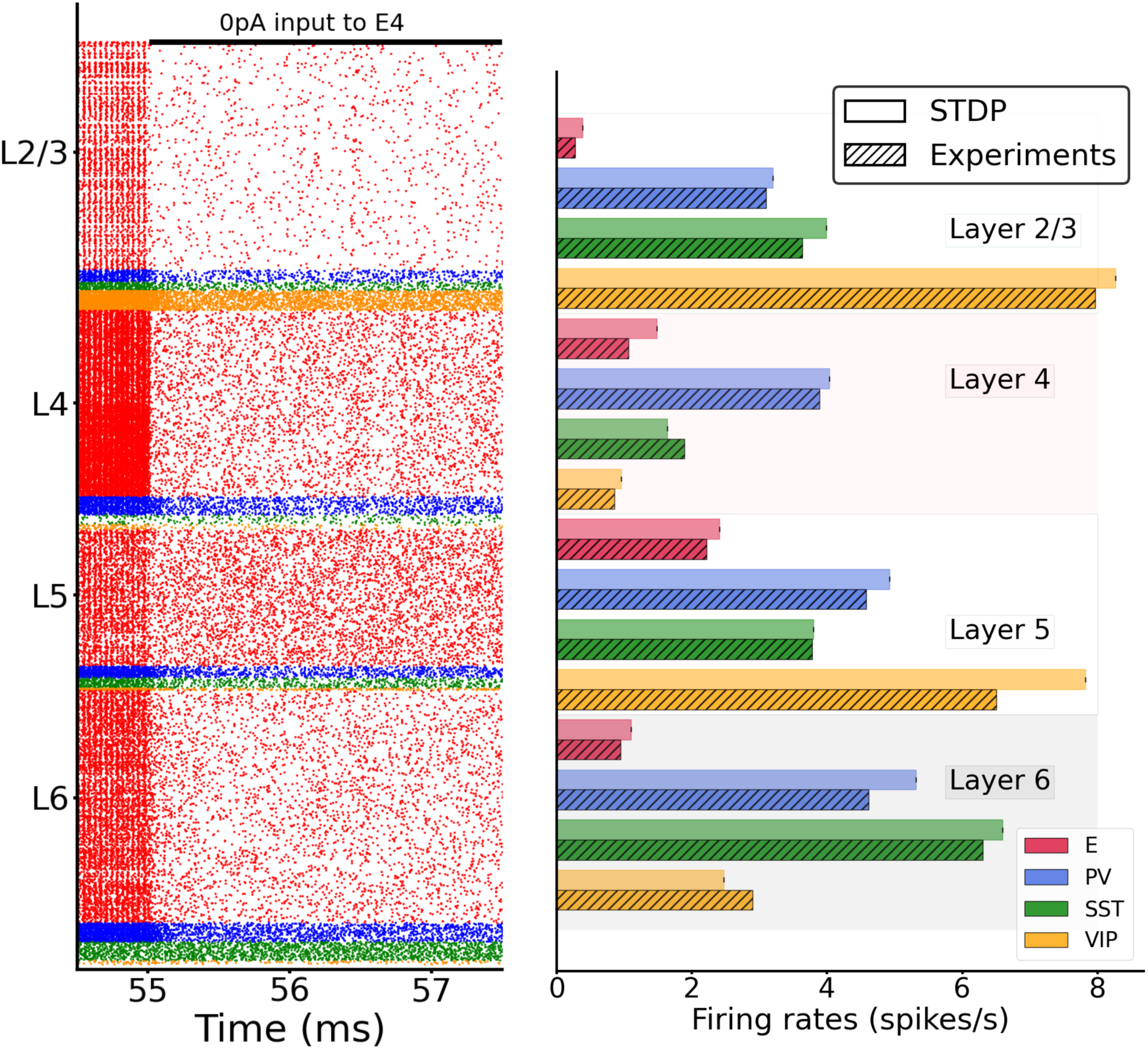
Spontaneous activity after STDP-induced column. Left: raster plot of spontaneous spiking activity simulated for 1500 ms after plasticity offset (55 s). Right: Mean firing rates for each model population versus experiment.

**Figure S5:**
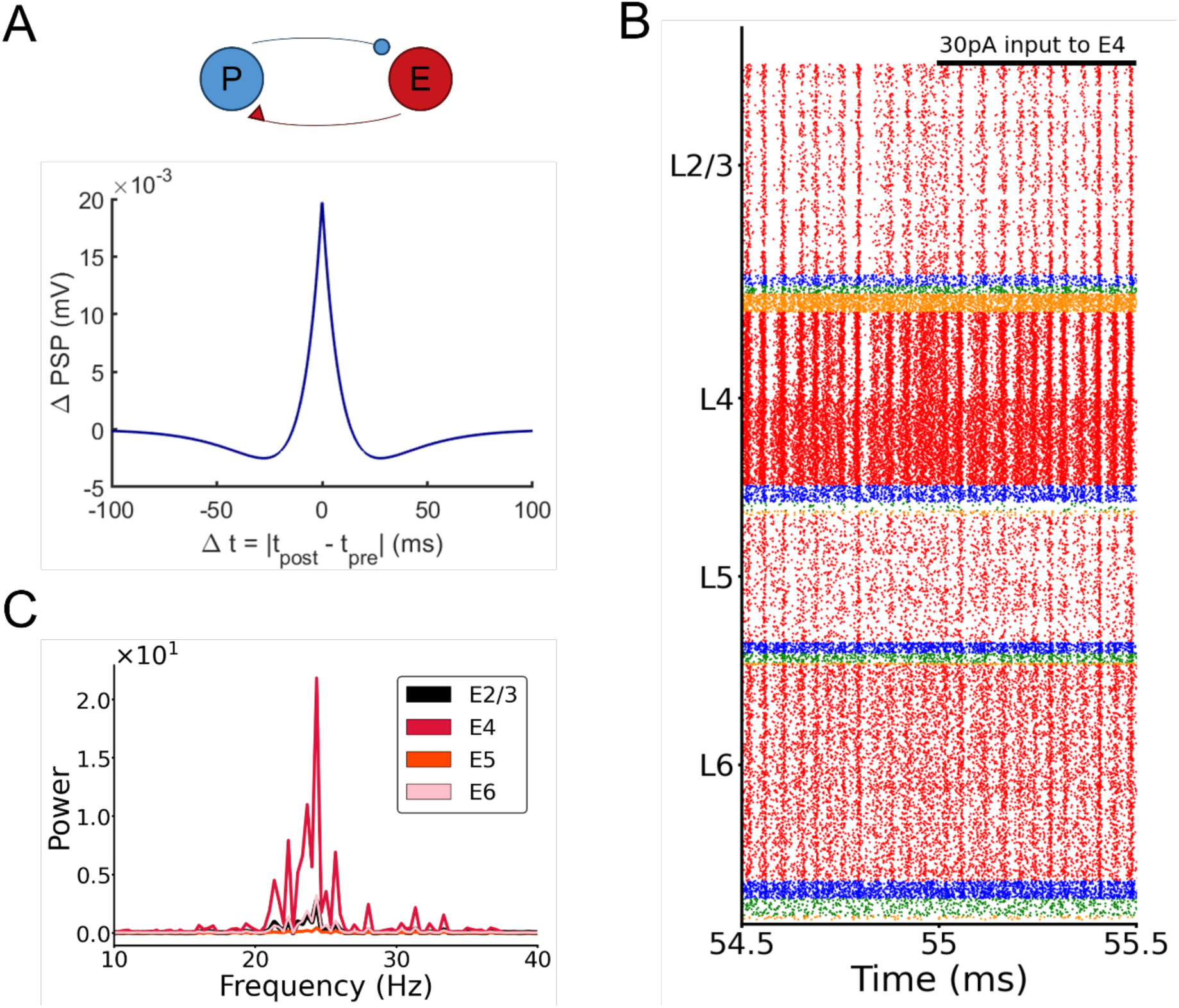
Emergence of gamma oscillations in a model with plastic PV-E weights in addition to the plastic E-E weights. (A) Scheme of the inhibitory STDP rule: the relationship between weight changes and relative timing between pre and postsynaptic spikes follows a symmetric “sunken Mexican hat” curve. (B) Raster plot of spike activity in the column with both excitatory and inhibitory plasticity, showing the emergence of oscillations. A continuous input of 30 pA is given to half of the pyramidal cells in layer 4. All excitatory-to-excitatory, as well as all PV-to-excitatory connections in the entire column evolve according to their corresponding STDP rule. (C) Power spectrum of the firing rates of pyramidal cells for different layers, showing a broadening of the curves and a decrease of peak power compared to the results with only excitatory plasticity (Fig. 4C).

**Figure S6:**
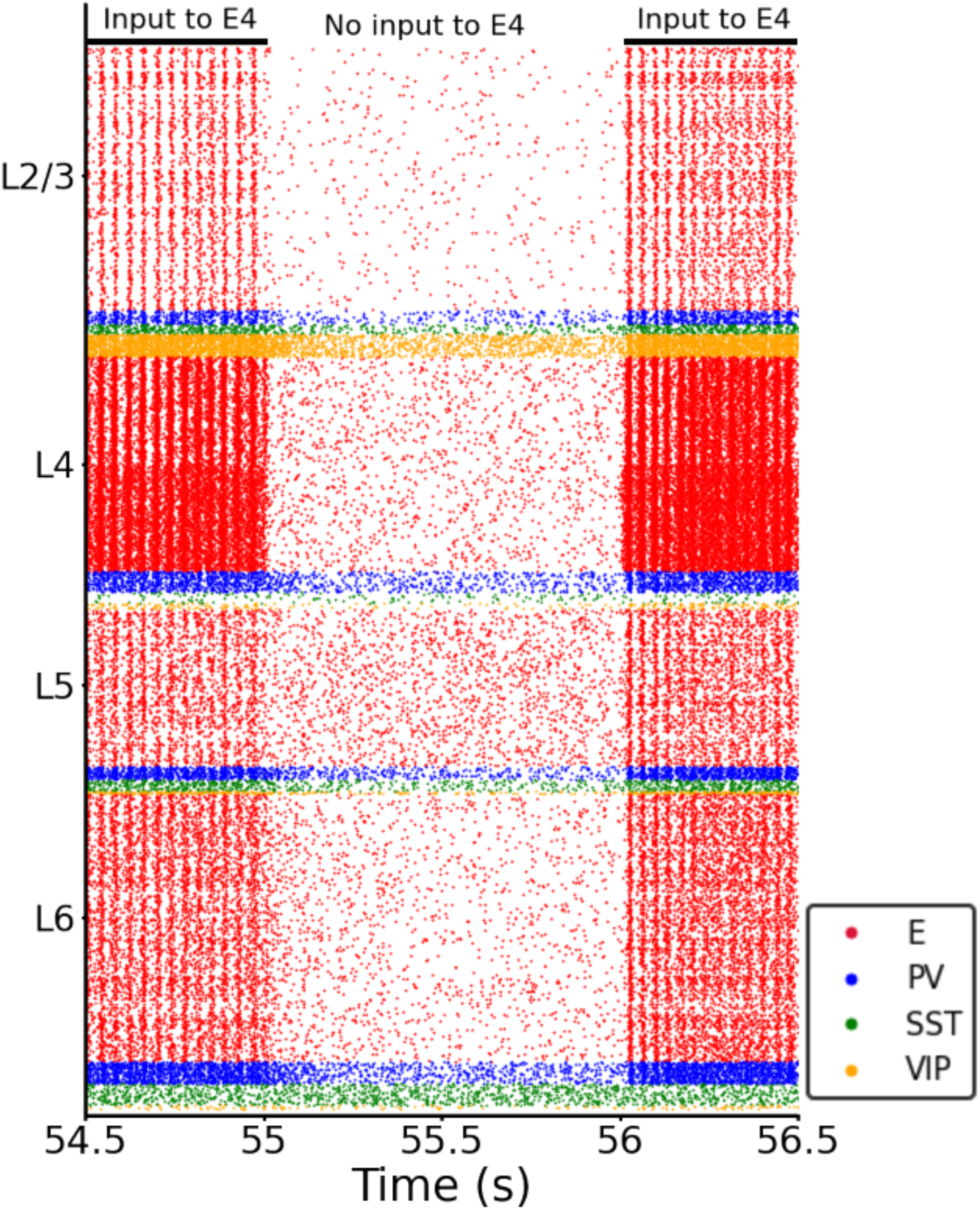
Raster plot showing the relations between input to layer 4 and oscillations. STDP plasticity was present from the start of the simulation until 55 s. When the input was switched off at t=55 s, the oscillations (frequency: 26 Hz) disappeared; vice versa for switching the input back on. All hyperparameters of this simulations are listed in Tables S3-S10.

**Figure S7:**
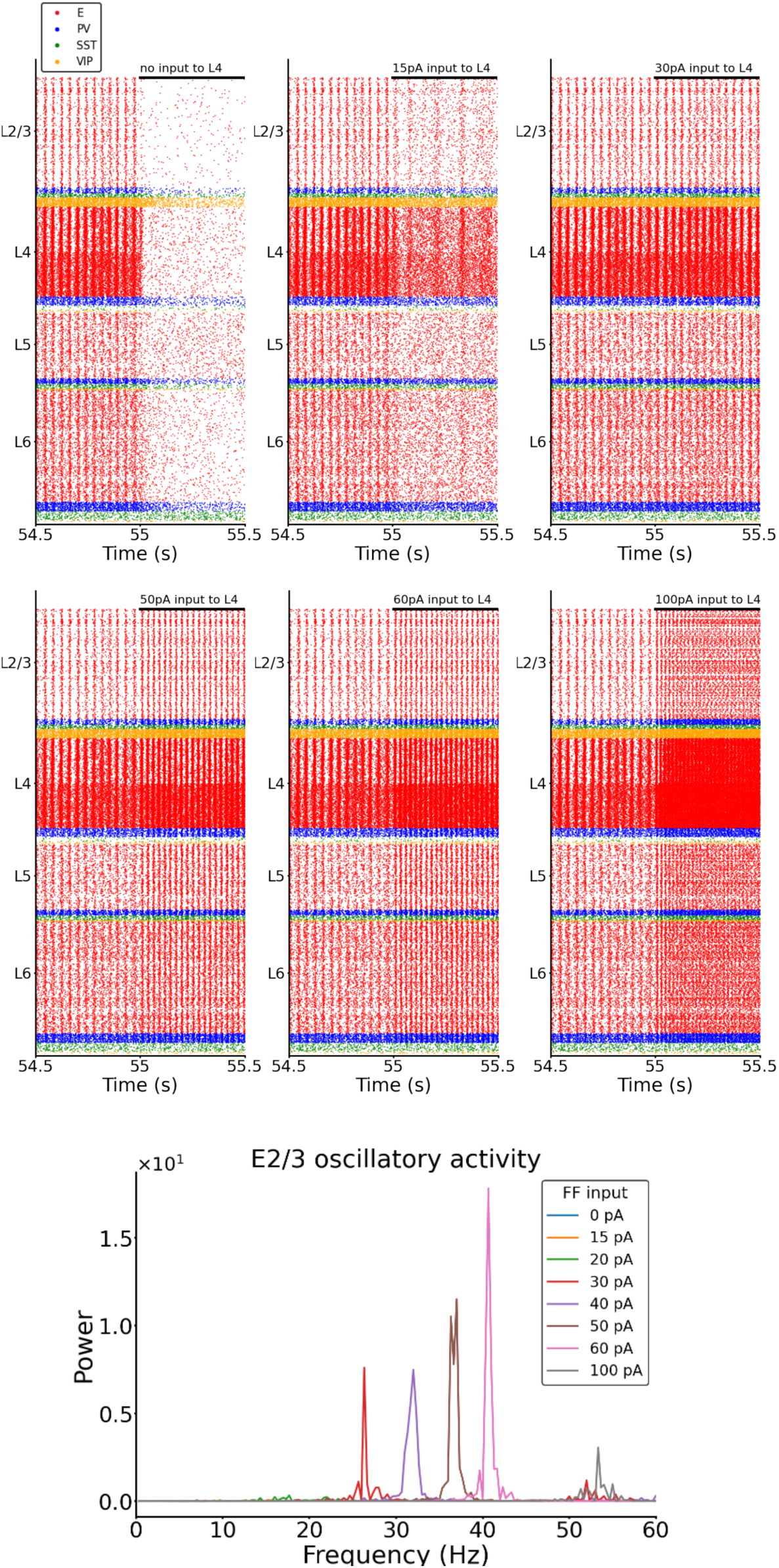
Modulation of oscillations by feedforward input strength. Top: Raster plots of a subset of the conditions shown in Fig.5A (varying input strength to excitatory cells in layer 4: no input, 15 pA, 30 pA, 50 pA, 60 pA, 100 pA) . Applying more input to layer 4 causes faster oscillations. When no input to layer 4 is present the oscillations disappear. Bottom: Power spectrum of the frequency of excitatory Layer 2/3 neuron firing with varying feedforward input to layer 4 (each color trace represents a different amount of feedforward input to layer 4). The peak amplitudes (max power) and the corresponding frequency are also shown in Fig. 5B and 5C.

**Figure S8:**
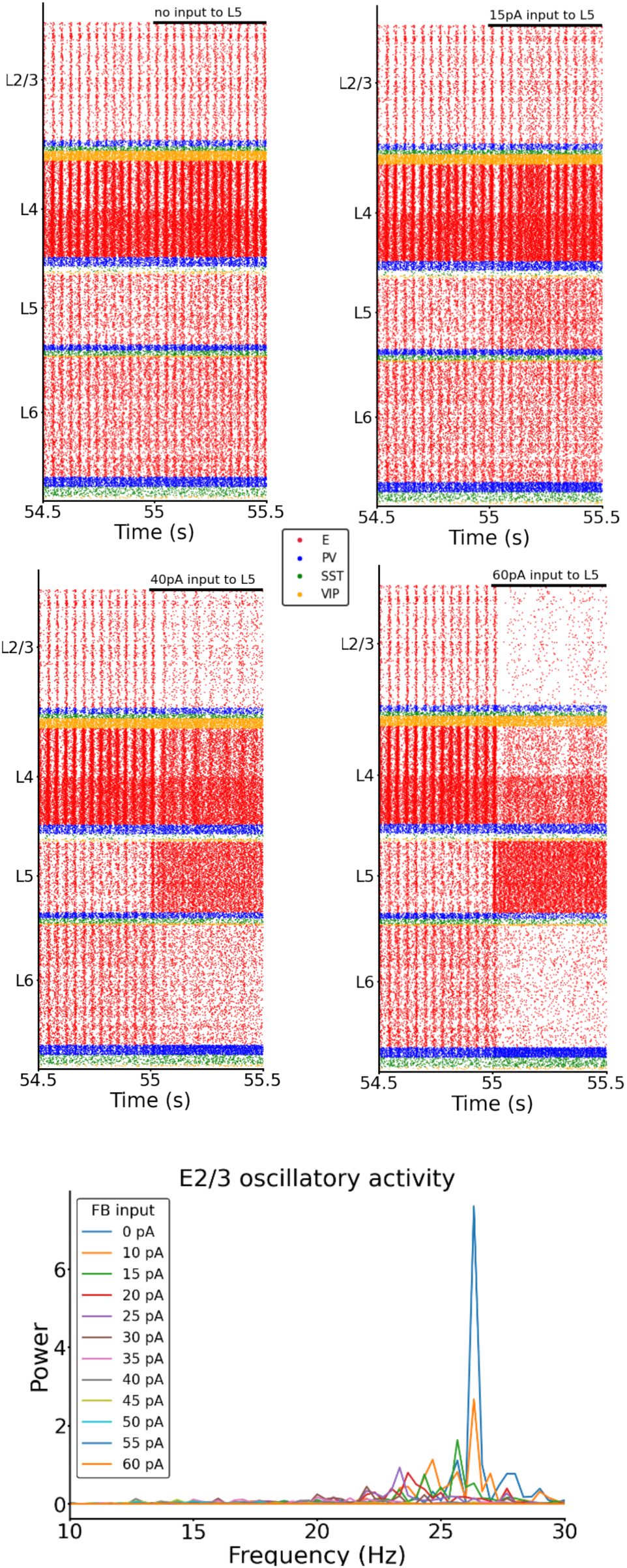
Modulation of oscillations by feedback (FB) input strength. Top: Raster plots of a subset of the conditions shown in Fig. 5D (varying input strength to excitatory cells in layer 5: no feedback input, 15 pA, 40 pA, 60pA) . The more input is applied to layer 5 (while keeping a constant input of 30 pA to layer 4) the more the oscillations slow down and decrease in power. Bottom: Power spectrum of the frequency of excitatory layer 2/3 neuron firing with varying feedback (FB) input (each color trace represents a different amount of feedback input to layer 5). The peak amplitudes (max power) and the corresponding frequency are also shown in Fig. 5E and 5F.

**Figure S9:**
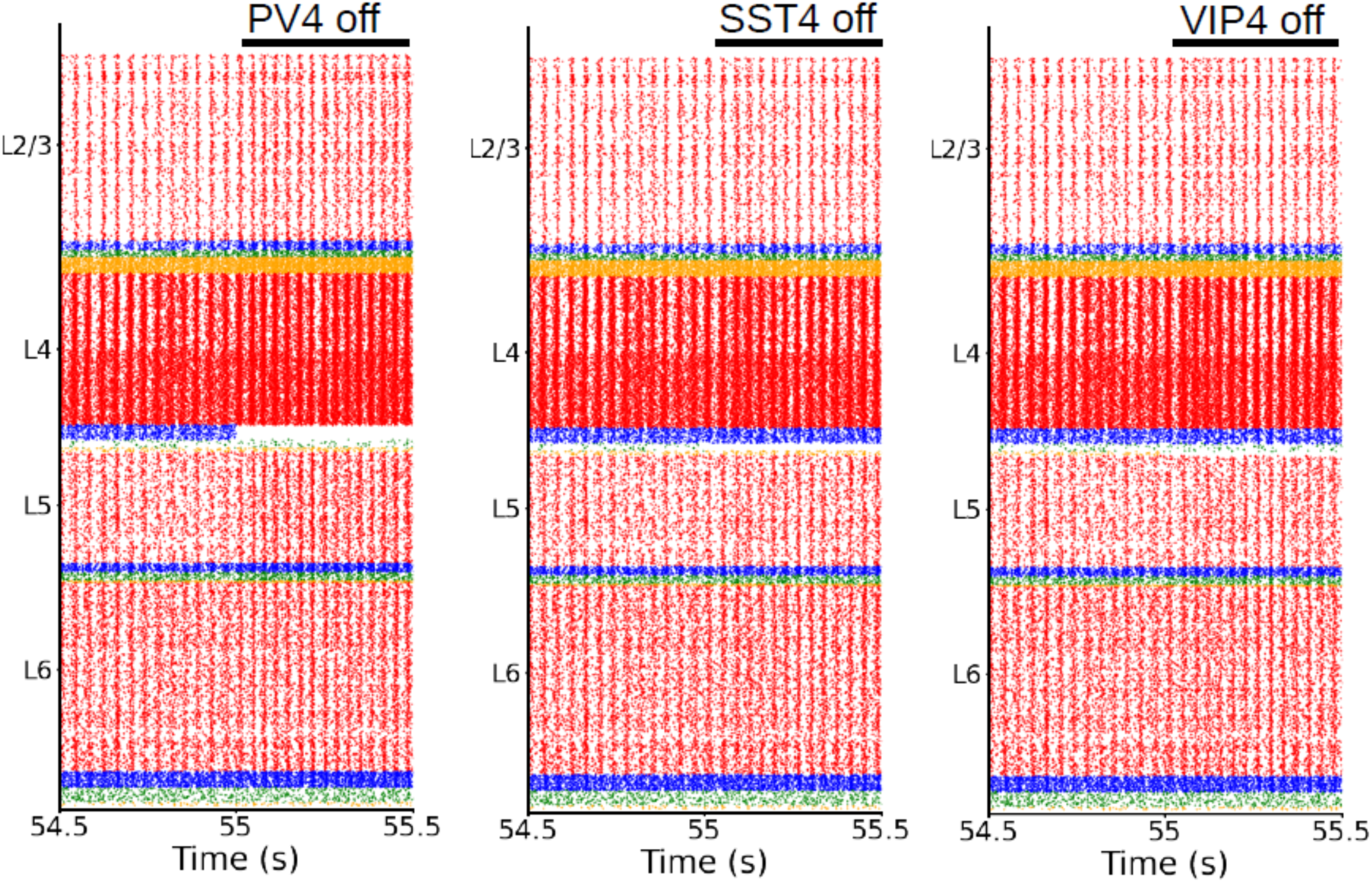
Raster plots of the whole column model for inactivating different cell groups in layer 4. Each group is inactivated (one at a time) at 55 s to visualize the effects on neural dynamics. From left to right: inactivation of PV neuron in layer 4, SST neurons in layer 4, VIP neurons in layer 4. Removing one group of inhibitory neurons at a time has only a minor effect on the oscillations: a slight increase of the oscillations speed. A bigger effect in increasing the speed of oscillations is however shown in the case of silencing PV cells (leftmost raster plot).

**Figure S10:**
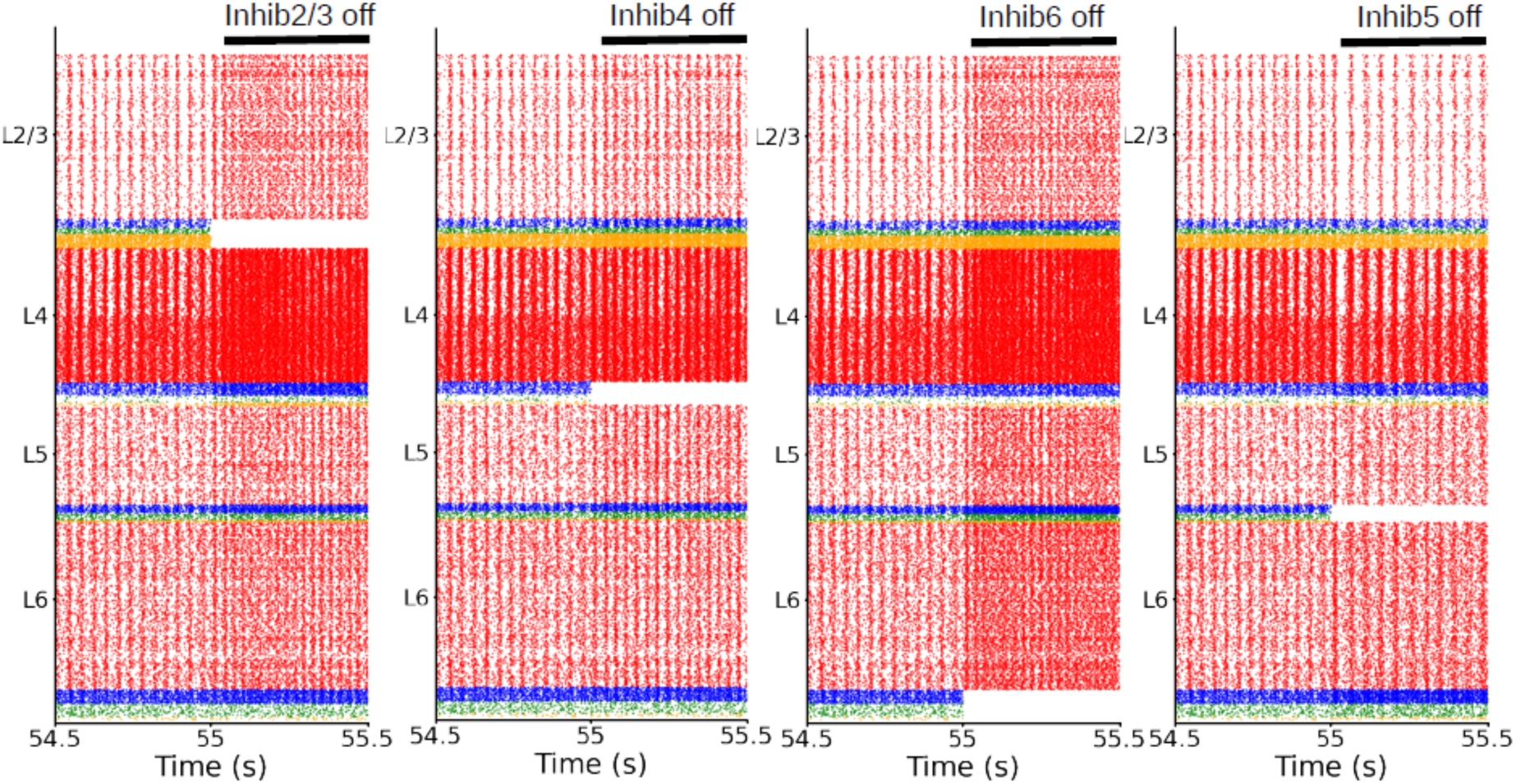
Raster plots of the whole column model for different inactivation conditions (layer analysis). All inhibitory cells in a layer are inactivated at 55 s, to visualize the effects on neural dynamics. From left to right: inactivation of all inhibitory neurons in layer 2/3, layer 4, layer 6, layer 5. All inhibitory neuron types in each layer contribute to oscillations, removing them one layer at a time shows their effect (increase of oscillation frequency). The inactivation of the inhibitory neurons in layer 2/3 shows the biggest effect. Layer 5 is the only exception: here the oscillations decrease in frequency and power.

**Figure S11:**
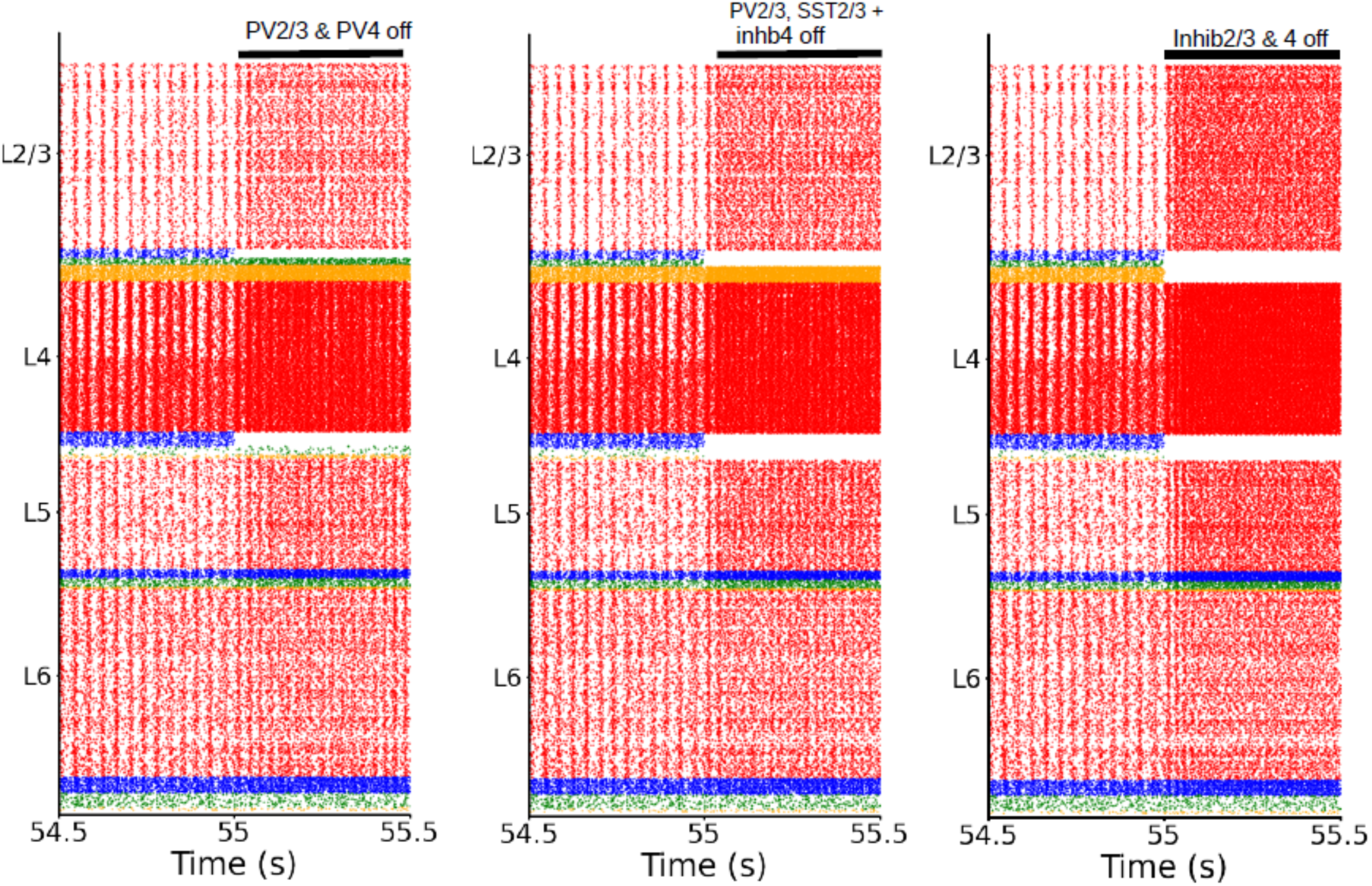
Raster plots of the whole column model for different inactivation conditions. Each combination of groups is inactivated at 55 s. From left to right: inactivation of PV cells in layer 2/3 and PV cells in layer 4; inactivation of PV and SST cells in layer 2/3; and all inhibitory cells in layer 4, inactivation of all inhibitory neurons in layer 2/3 and 4. Removing more and more groups (from left to right) causes a significant increase in oscillation frequency.

**Figure S12:**
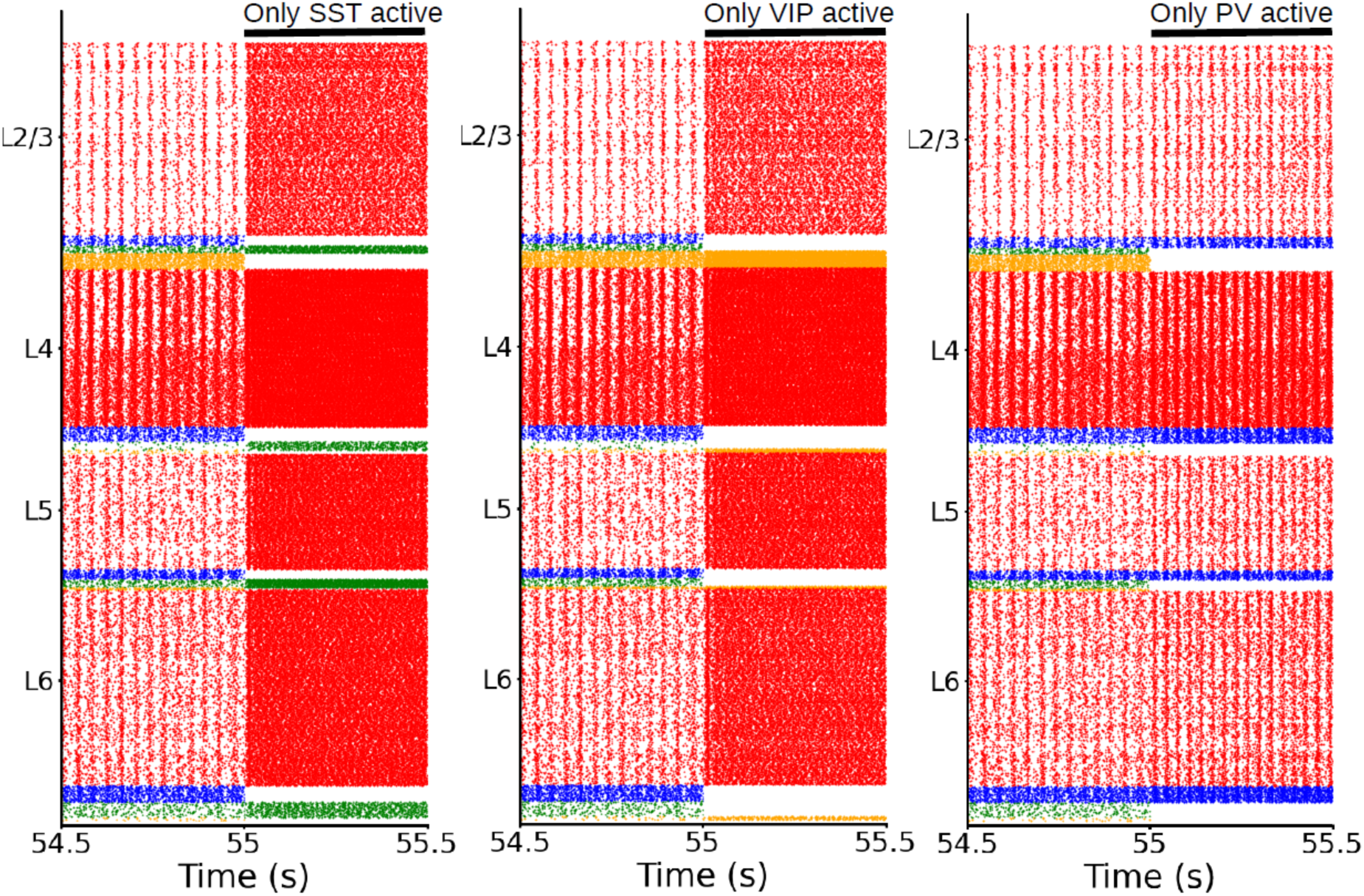
Raster plots of the whole column model for different inactivation conditions. Each cell type across all layers is inactivated at 55 s. These are experiments where we are leaving active only one type of inhibitory neurons in the entire column (next to pyramidal cells). From left to right: only SST active, only VIP active, only PV active. This shows that – together with pyramidal cells – PV cells alone are able to maintain oscillations around the same frequency (26 Hz). Only a slight increase is visible, the same is not true for SST or VIP. When only them are active the oscillations are drastically affected (first two raster plots).

**Figure S13:**
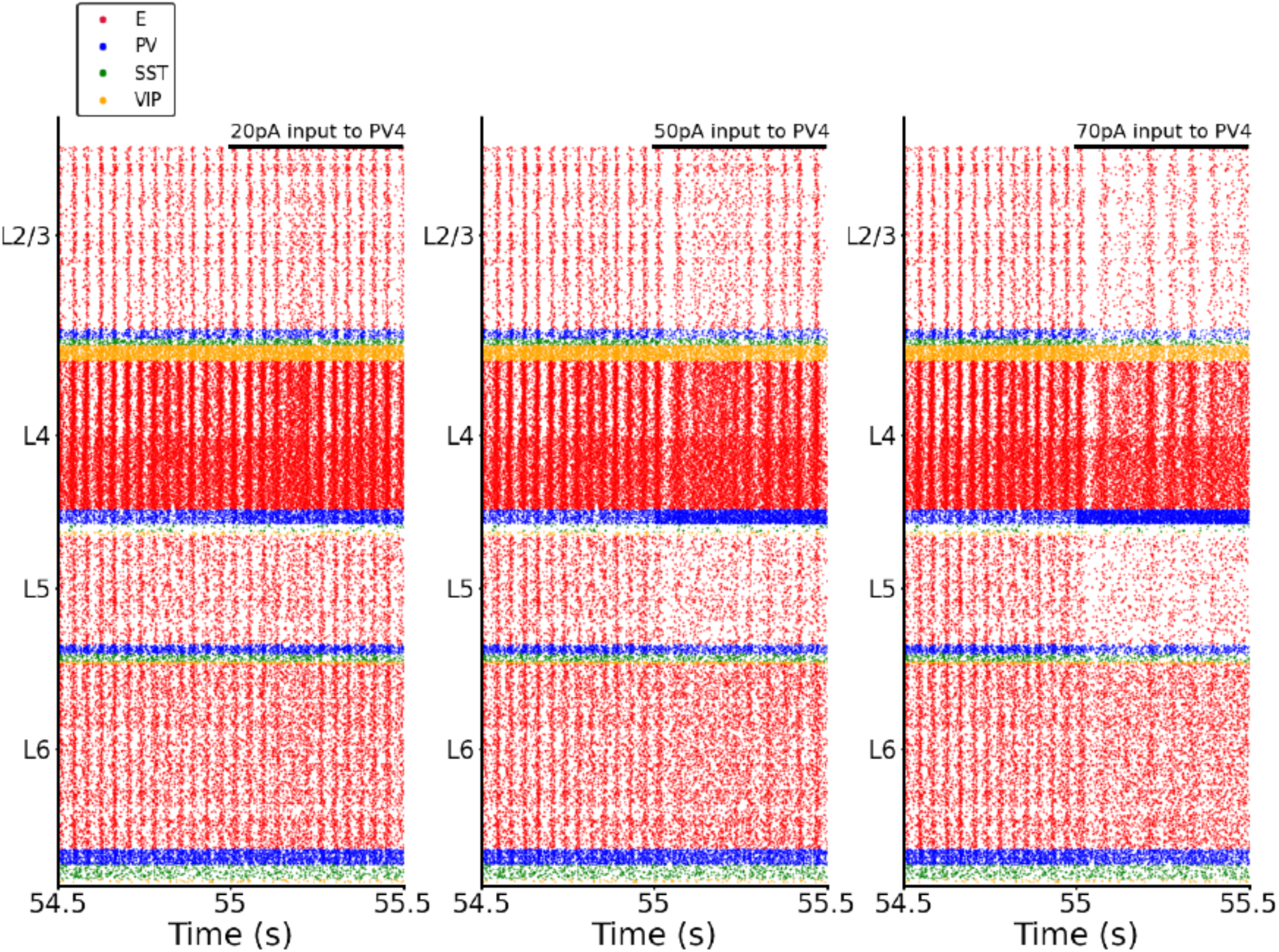
Raster plots of the whole column model for different input strengths applied to PV neurons in layer 4. The input is given at 55 s. From left to right: 20 pA of input to PV in layer 4, 50 pA of input to PV in layer 4, 70 pA of input to PV in layer 4. Injecting more input into PV cells shows that they are able to modulate the frequency of the oscillations. The more input to layer 4 PV the more the oscillation frequency decreases. In the first two raster plots, a blurring of the oscillations can be observed after 55 seconds. The excitatory neurons are receiving increased inhibition from the PV cells (which are also receiving external input). As a result, the synchrony of their firing, which generates the oscillations, is partially lost, leading to this observable effect.

**Figure S14:**
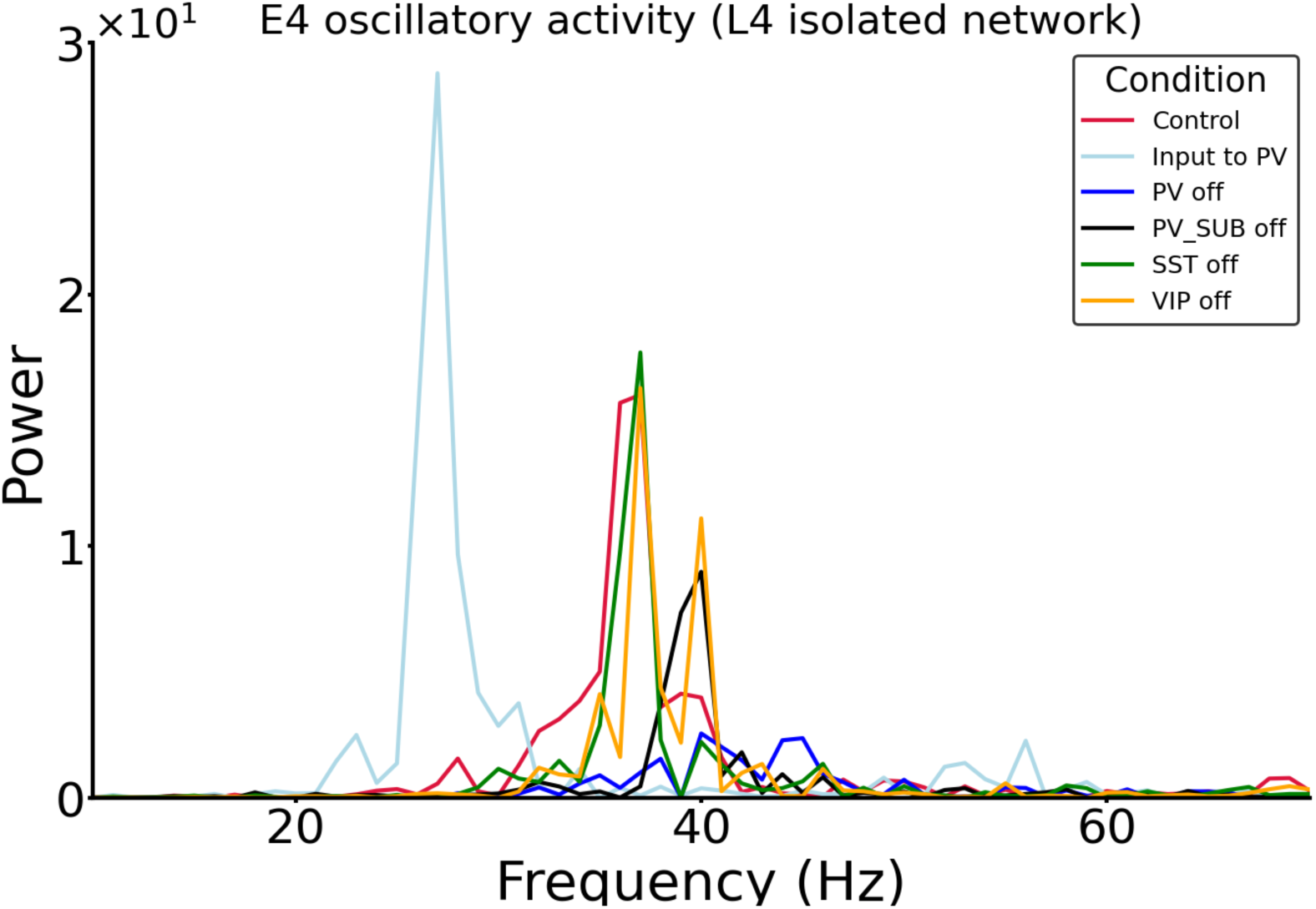
Power spectrum of the frequency of excitatory neuron firing in layer 4 for different inactivation conditions for the isolated network of layer 4 (schematics in Fig. 6C left). Each color trace represents a different condition, in the legend the name of the group (PV, SST, VIP) represents the inhibited group. Here we can appreciate the shift in the power peak depending on the analyzed condition: when PV cells are inactivated the frequency with max power (peak in the plot) is shifted to the right (blue), indicating an increase in speed of the oscillations. In contrast, when PV cells are stimulated by an external input, they modulate the frequency of the oscillations which shows a significant decrease (light blue trace: Input to PV). This result is consistent with the full-model scenario shown in Fig. S13. A total inhibition of SST cells or VIP cells is not significantly affecting the oscillations (green and yellow trace). The maxima of oscillatory power at 37 Hz for the different conditions were also shown in Fig. 6C (left) as well as the frequency where maximal power was reached (right in Fig. 6C).

**Figure S15:**
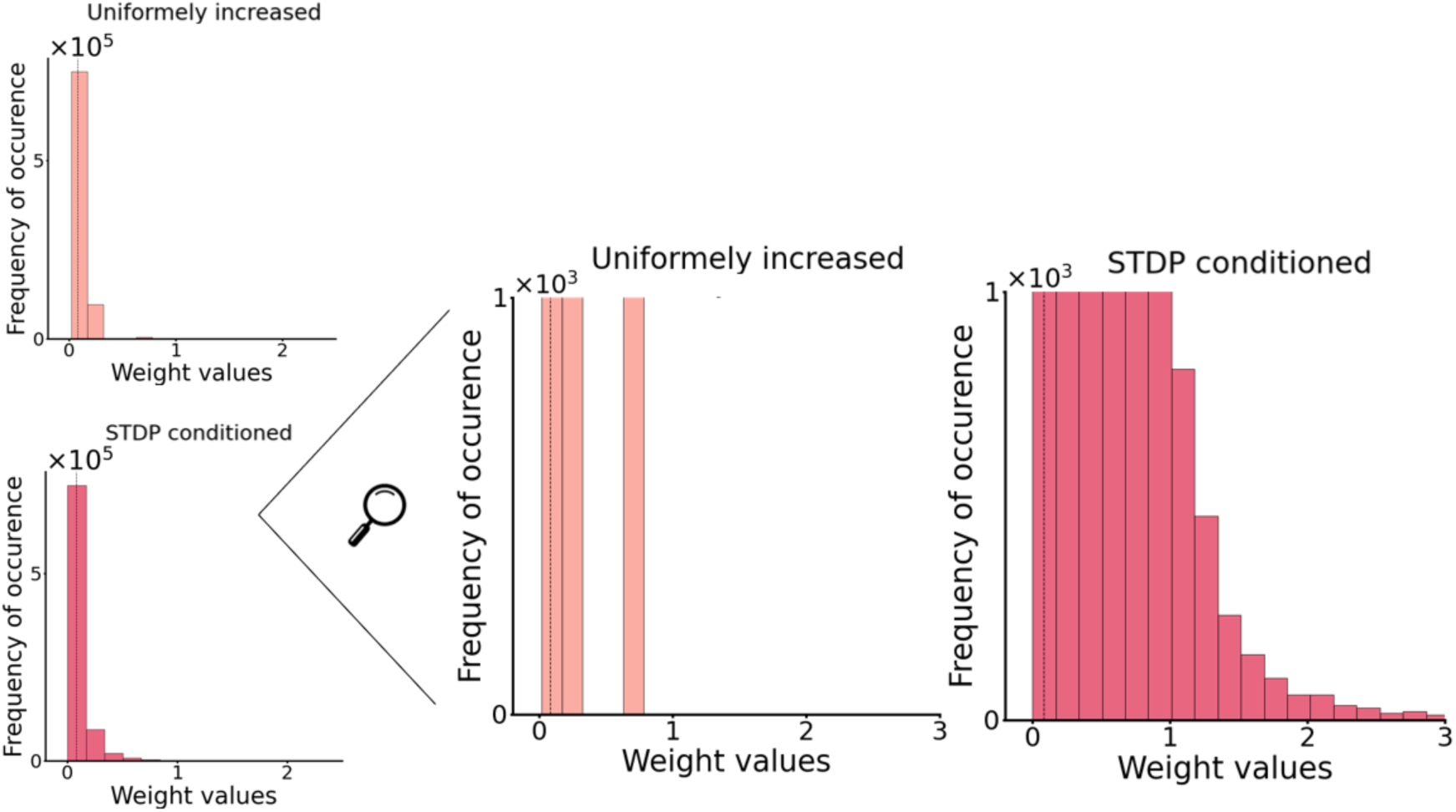
Weighs distribution for the trained network and uniformly increased network (see Fig. 7). Left: distribution of weights for the two networks. Right: zoom in by cutting the threshold on the y-axis at 10^!^to better visualize the differences between the distributions. Even if the average of all connection strengths of the STDP conditioned and uniformly increased network are the same (mean=0.081) a closer look at the left panel shows that the distributions are different. In the Uniformly increased network, there are few connections (less than 600 out of the total of 800k) that have a high value ( >1.5). Those strong connections are absent in the Uniformly increased network. The reason that the means of the two distributions are so similar, is that in the STDP conditioned network the vast majority of connections are very small (∼0), which collectively compensates for the small subset of strong connections. In the uniformly increased network those strong connections are absent, in conjunction with the absence oscillations.

**Figure S16:**
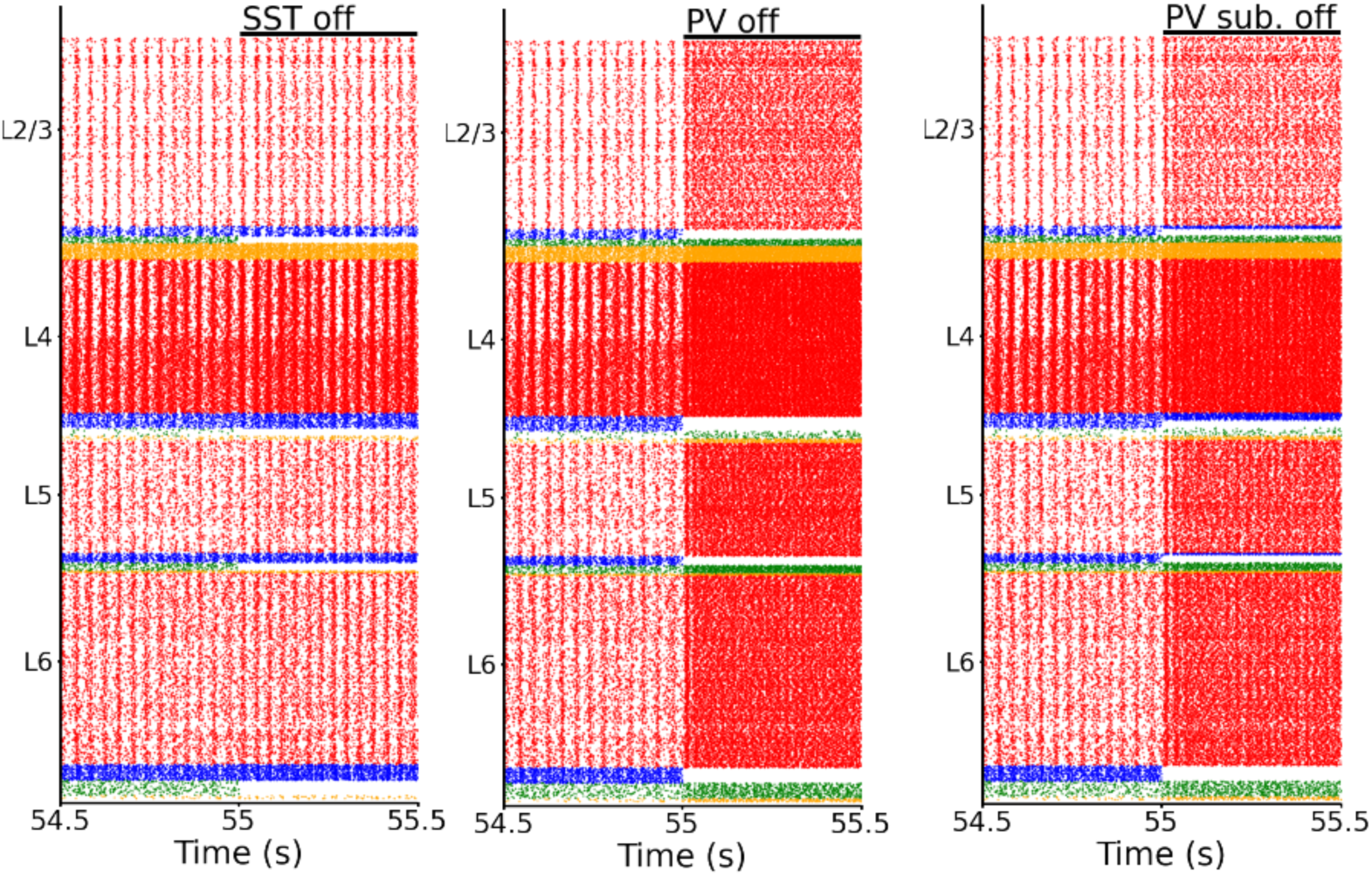
Raster plots of the entire column model under different inactivation conditions. Each cell type across all layers is inactivated at 55 seconds. The first two raster plots are also shown in 6B. From left to right: the effect of silencing all SST cells, all PV cells, and a subset of PV cells. To test whether the impact of PV cells was not solely due to the higher number of PV cells compared to SST cells, we inactivated a subpopulation of PV cells (PV-sub), equal in size to the number of SST neurons in each layer. The effects on the oscillations, in this case, were significantly stronger than the inactivation of the SST population, thereby proving the stronger role of PV cells. The prominent role of PV cells appeared to be related, not solely to their higher number, but rather to the synaptic connections from PV to pyramidal neurons, which are stronger than the projections from SST to pyramidal neurons.

**Figure S17:**
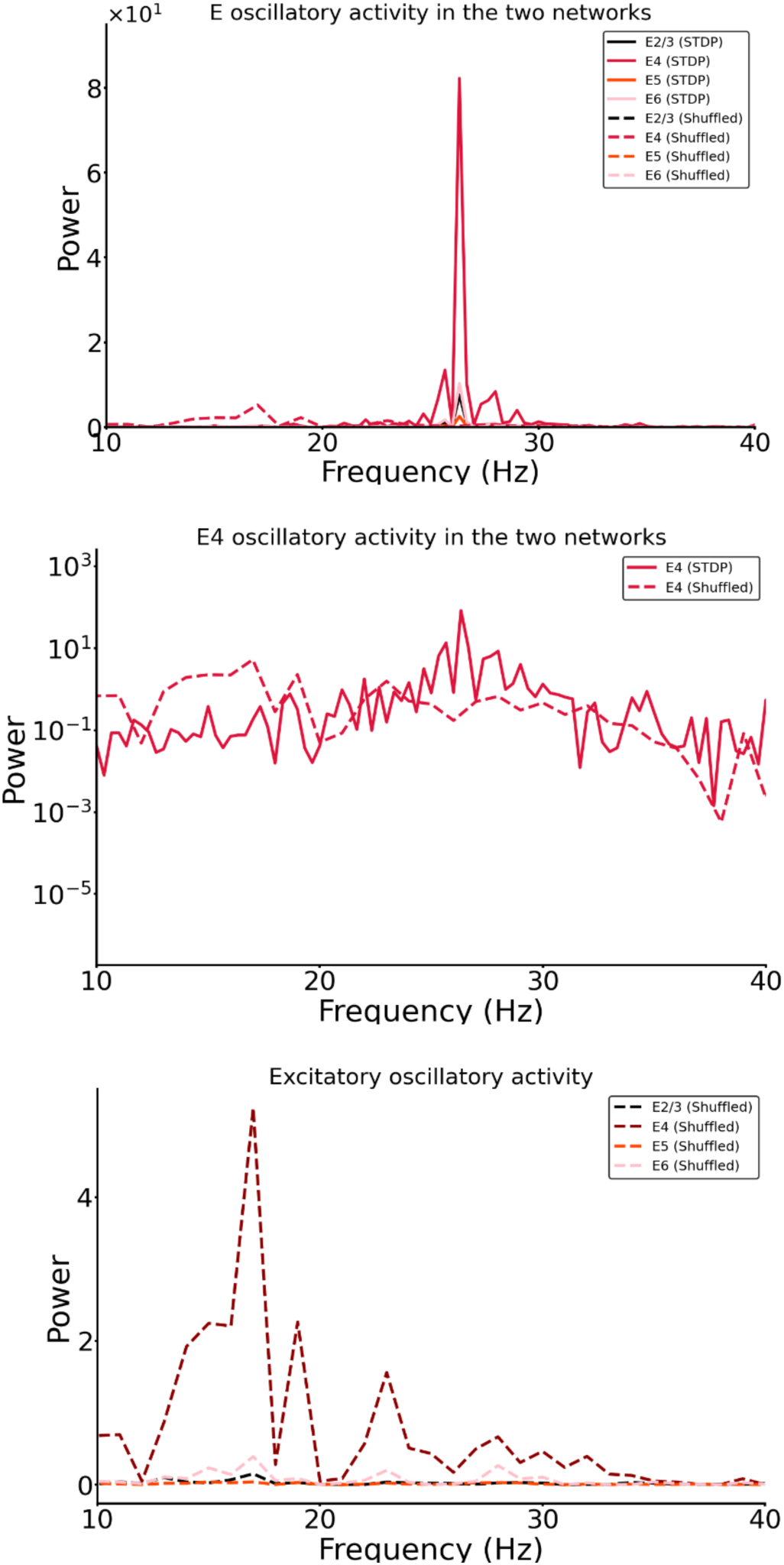
Top: Power spectrum of the frequency of excitatory neuron firing rates in all layers of two distinct networks (STDP conditioned and Shuffled networks). As shown in Figure 7A, in the network with shuffled weights, the oscillatory activity disappears. The peaks at 26 Hz in the Shuffled network are no longer visible. Middle: Power spectrum of the frequency of excitatory neuron firing rates in layer 4 of the two distinct networks (STDP conditioned and Shuffled networks). A logarithmic scale is used to better appreciate the power drop. Bottom: Power spectrum of the frequency of excitatory neuron firing rates in all layers for the Shuffled network. The maximum peak is at 17 Hz; the power at 26 Hz is now irrelevant compared to the STDP conditioned network. In the Shuffled network, the power at 26 Hz drastically drops (e.g., value drops from the scale 10^0^to 10^−3^ for excitatory neurons in Layer 2/3), see also Table 10.

## Tables

**Table 1:** Number of neurons in each layer. N_tot_ is the total number of cells in the column and can be defined arbitrarily for a simulation. The number of cells in each layer will scale accordingly. We used N_tot_=5000 for all main simulations. In S2 we showed the results for N_tot_=10000 and N_tot_=20000.

|  | Number of neurons |
| --- | --- |
| L1 | $0.0192574218 \cdot N_{\text{tot}}$ |
| L2/3 | $0.291088453 \cdot N_{\text{tot}}$ |
| L4 | $0.237625904 \cdot N_{\text{tot}}$ |
| L5 | $0.17425693 \cdot N_{\text{tot}}$ |
| L6 | $0.297031276 \cdot N_{\text{tot}}$ |

**Table 2:** Percentage of inhibitory neurons as a fraction of the total number of inhibitory cells in each layer. In each layer, the inhibitory cells represent 15% of the total number of neurons for that layer.

| Percentage (%) | PV | SST | VIP |
| --- | --- | --- | --- |
| L2/3 | 0.295918 | 0.214286 | 0.489796 |
| L4 | 0.552381 | 0.295238 | 0.152381 |
| L5 | 0.485714 | 0.428571 | 0.085714 |
| L6 | 0.458333 | 0.458333 | 0.083333 |

**Table 3:** Cell counts in the network for N_tot_ = 5000.

| Number of neurons | E | PV | SST | VIP |
| --- | --- | --- | --- | --- |
| L1 |  |  |  | 96 |
| L2/3 | 1236 | 65 | 47 | 107 |
| L4 | 1010 | 98 | 53 | 27 |
| L5 | 741 | 63 | 56 | 11 |
| L6 | 1263 | 102 | 102 | 19 |

**Table 4:**
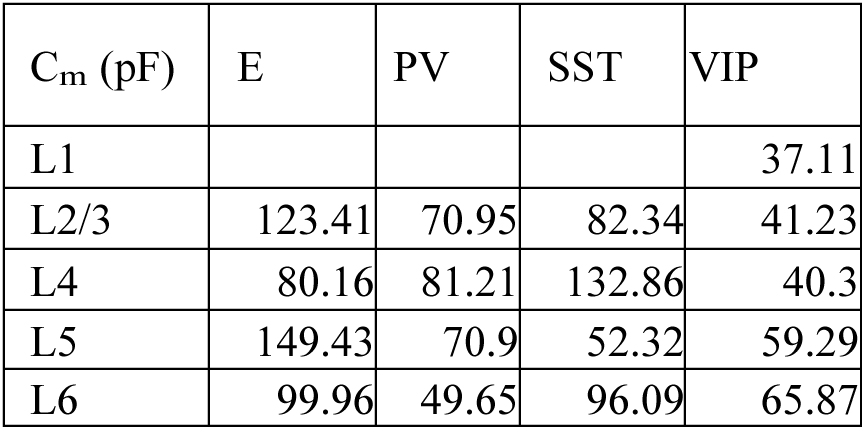
Membrane capacitance for each group of cells. The values are taken from the Allen database^22^. The same values are used for all simulations.

| $C_m$ (pF) | E | PV | SST | VIP |
| --- | --- | --- | --- | --- |
| L1 |  |  |  | 37.11 |
| L2/3 | 123.41 | 70.95 | 82.34 | 41.23 |
| L4 | 80.16 | 81.21 | 132.86 | 40.3 |
| L5 | 149.43 | 70.9 | 52.32 | 59.29 |
| L6 | 99.96 | 49.65 | 96.09 | 65.87 |

**Table 5:** Leak conductance for each group of cells^22^.

| $gL$ (nS) | E | PV | SST | VIP |
| --- | --- | --- | --- | --- |
| L1 |  |  |  | 4.07 |
| L2/3 | 2.47 | 9.49 | 3.17 | 6.4 |
| L4 | 5.16 | 9.19 | 7.96 | 1.87 |
| L5 | 16.66 | 5.21 | 3.43 | 6.52 |
| L6 | 5.88 | 6.86 | 2.99 | 6.09 |

**Table 6:**
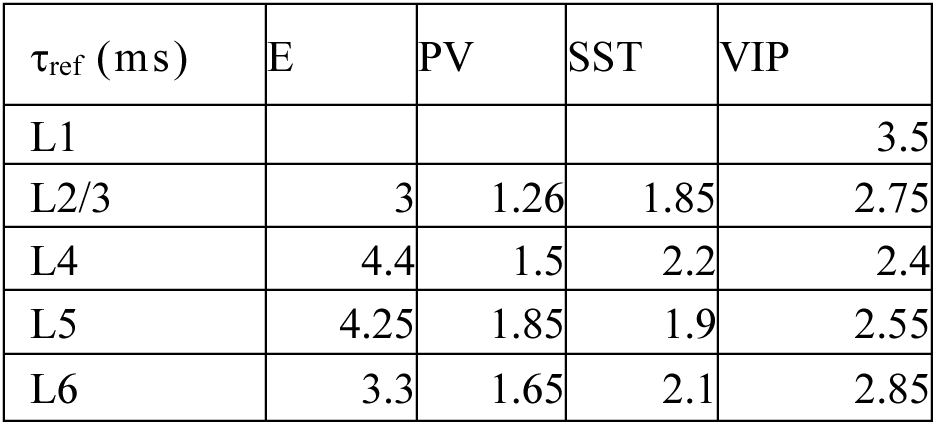
Refractory period for each group of cells^22^.

| $\tau_{\text{ref}}$ (ms) | E | PV | SST | VIP |
| --- | --- | --- | --- | --- |
| L1 |  |  |  | 3.5 |
| L2/3 | 3 | 1.26 | 1.85 | 2.75 |
| L4 | 4.4 | 1.5 | 2.2 | 2.4 |
| L5 | 4.25 | 1.85 | 1.9 | 2.55 |
| L6 | 3.3 | 1.65 | 2.1 | 2.85 |

**Table 7:** Resting membrane potential for each group of cells^22^.

| $V_{\text{rest}}$ (mV) | E | PV | SST | VIP |
| --- | --- | --- | --- | --- |
| L1 |  |  |  | -65.5 |
| L2/3 | -80.97 | -82.35 | -69.16 | -67.94 |
| L4 | -72.53 | -70.45 | -74.2 | -63.14 |
| L5 | -68.28 | -77.5 | -70.01 | -72.00 |
| L6 | -77.5 | -76.42 | -62.99 | -78.85 |

**Table 8:** Spike threshold for each group of cells^22^.

| $V_{\text{th}}$ (mV) | E | PV | SST | VIP |
| --- | --- | --- | --- | --- |
| L1 |  |  |  | -40.20 |
| L2/3 | -40.53 | -56.32 | -39.95 | -41.34 |
| L4 | -47.63 | -44.23 | -44.07 | -40.89 |
| L5 | -40.55 | -51.2 | -47.38 | -51.2 |
| L6 | -42.31 | -49.06 | -37.19 | -44.81 |

**Table 9:**
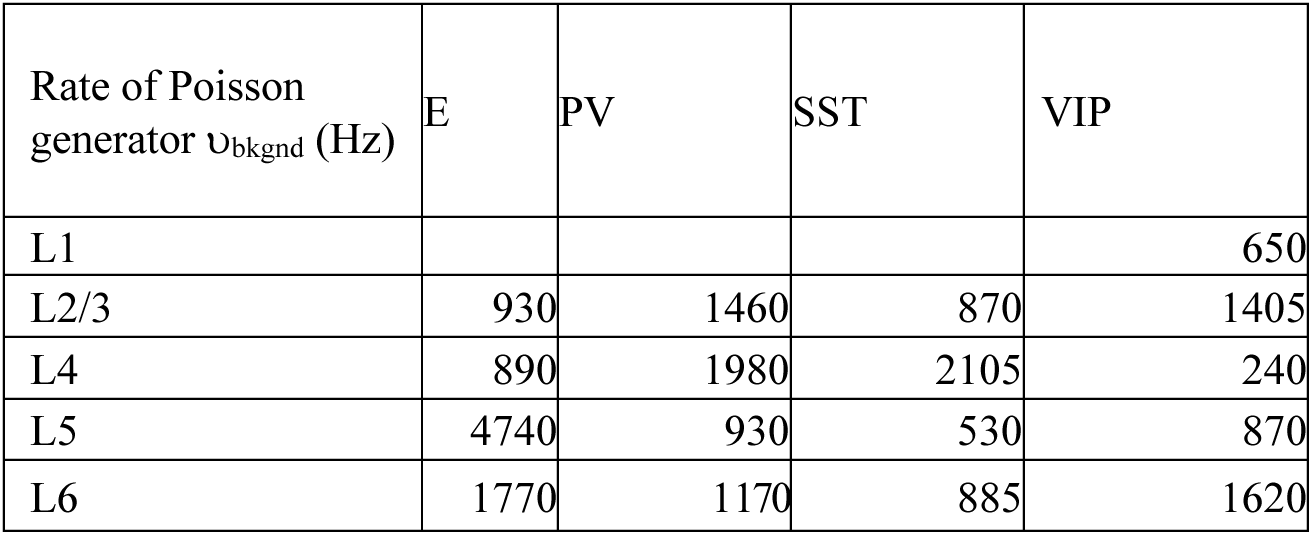
Background noise level that each group is receiving. Values represent the Poisson generator rates υ_bkgnd_. Synapses transmitting this external input to the neurons in the column have a synaptic strength of w_ext_ =1.

| Rate of Poisson generator $\nu_{bkgnd}$ (Hz) | E | PV | SST | VIP |
| --- | --- | --- | --- | --- |
| L1 |  |  |  | 650 |
| L2/3 | 930 | 1460 | 870 | 1405 |
| L4 | 890 | 1980 | 2105 | 240 |
| L5 | 4740 | 930 | 530 | 870 |
| L6 | 1770 | 1170 | 885 | 1620 |

**Table 10:** Peak power spectrum value of the excitatory neuron firing rates in each layer, for STDP-conditioned networks, shuffled networks at 26 Hz (frequency with maximum power for the STDP conditioned network) and 17 Hz (frequency with maximum power for the Shuffled network). Columns 3-6 correspond to the values of the power spectrum for pyramidal firing rate (2/3, 4, 5, 6) for each of the three cases above and for the frequencies specified in column 2. See also Figure S17.

| Network type | Frequency (Hz) | E2/3 | E4 | E5 | E6 |
| --- | --- | --- | --- | --- | --- |
| STDP conditioned | 26 | $7.6 * 10^0$ | $8.2 * 10^1$ | $2.6 * 10^0$ | $1.0 * 10^1$ |
| Shuffled | 26 | $7.7 * 10^{-3}$ | $7.8 * 10^{-4}$ | $7.2 * 10^{-4}$ | $4.7 * 10^{-3}$ |
| Shuffled | 17 | $1.5 * 10^{-1}$ | $5.2 * 10^0$ | $3.7 * 10^{-2}$ | $3.9 * 10^{-1}$ |

## References

Albada, S. J. van, Helias, M., and Diesmann, M. (2015). Scalability of Asynchronous Networks Is Limited by One-to-One Mapping between Effective Connectivity and Correlations. PLOS Comput. Biol. 11, e1004490. doi: 10.1371/journal.pcbi.1004490

Antonoudiou, P., Tan, Y. L., Kontou, G., Upton, A. L., and Mann, E. O. (2020). Parvalbumin and Somatostatin Interneurons Contribute to the Generation of Hippocampal Gamma Oscillations. J. Neurosci. 40, 7668–7687. doi: 10.1523/JNEUROSCI.0261-20.2020

Bastos, A. M., Vezoli, J., Bosman, C. A., Schoffelen, J.-M., Oostenveld, R., Dowdall, J. R., et al. (2015). Visual Areas Exert Feedforward and Feedback Influences through Distinct Frequency Channels. Neuron 85, 390–401. doi: 10.1016/j.neuron.2014.12.018

Battaglia, D., and Hansel, D. (2011). Synchronous Chaos and Broad Band Gamma Rhythm in a Minimal Multi-Layer Model of Primary Visual Cortex. PLOS Comput. Biol. 7, e1002176. doi: 10.1371/journal.pcbi.1002176

Beerendonk, L., Mejías, J. F., Nuiten, S. A., de Gee, J. W., Fahrenfort, J. J., and van Gaal, S. (2024). A disinhibitory circuit mechanism explains a general principle of peak performance during mid-level arousal. Proc. Natl. Acad. Sci. 121, e2312898121. doi: 10.1073/pnas.2312898121

Bernander, O., Douglas, R. J., Martin, K. A., and Koch, C. (1991). Synaptic background activity influences spatiotemporal integration in single pyramidal cells. Proc. Natl. Acad. Sci. 88, 11569–11573. doi: 10.1073/pnas.88.24.11569

Billeh, Y. N., Cai, B., Gratiy, S. L., Dai, K., Iyer, R., Gouwens, N. W., et al. (2020). Systematic Integration of Structural and Functional Data into Multi-scale Models of Mouse Primary Visual Cortex. Neuron 106, 388–403.e18. doi: 10.1016/j.neuron.2020.01.040

Binzegger, T., Douglas, R. J., and Martin, K. A. C. (2004). A Quantitative Map of the Circuit of Cat Primary Visual Cortex. J. Neurosci. 24, 8441–8453. doi: 10.1523/JNEUROSCI.1400-04.2004

Bosman, C. A., Lansink, C. S., and Pennartz, C. M. A. (2014). Functions of gamma-band synchronization in cognition: from single circuits to functional diversity across cortical and subcortical systems. Eur. J. Neurosci. 39, 1982–1999. doi: 10.1111/ejn.12606

Brunel, N. (2000). Dynamics of Sparsely Connected Networks of Excitatory and Inhibitory Spiking Neurons. J. Comput. Neurosci. 8, 183–208. doi: 10.1023/A:1008925309027

Brunel, N., and Wang, X.-J. (2003). What Determines the Frequency of Fast Network Oscillations With Irregular Neural Discharges? I. Synaptic Dynamics and Excitation-Inhibition Balance. J. Neurophysiol. 90, 415–430. doi: 10.1152/jn.01095.2002

Cardin, J. A., Carlén, M., Meletis, K., Knoblich, U., Zhang, F., Deisseroth, K., et al. (2009). Driving fast-spiking cells induces gamma rhythm and controls sensory responses. Nature 459, 663–667. doi: 10.1038/nature08002

Douglas, R. J., and Martin, K. A. C. (2004). Neuronal Circuits of the Neocortex. Annu. Rev. Neurosci. 27, 419–451. doi: 10.1146/annurev.neuro.27.070203.144152

Dowdall, J. R., and Vinck, M. (2023). Coherence fails to reliably capture inter-areal interactions in bidirectional neural systems with transmission delays. NeuroImage 271, 119998. doi: 10.1016/j.neuroimage.2023.119998

Feng, M., Bandyopadhyay, A., and Mejias, J. F. (2023). Emergence of distributed working memory in a human brain network model. 2023.01.26.525779. doi: 10.1101/2023.01.26.525779

Fries, P. (2005). A mechanism for cognitive dynamics: neuronal communication through neuronal coherence. Trends Cogn. Sci. 9, 474–480. doi: 10.1016/j.tics.2005.08.011

Garcia del Molino, L. C., Yang, G. R., Mejias, J. F., and Wang, X.-J. (2017). Paradoxical response reversal of top-down modulation in cortical circuits with three interneuron types. eLife 6, e29742. doi: 10.7554/eLife.29742

Gerstner, W., Kempter, R., van Hemmen, J. L., and Wagner, H. (1996). A neuronal learning rule for sub-millisecond temporal coding. Nature 383, 76–78. doi: 10.1038/383076a0

Gilbert, C. D. (1983). Microcircuitry of the Visual Cortex. Annu. Rev. Neurosci. 6, 217–247. doi: 10.1146/annurev.ne.06.030183.001245

Golomb, D. (2007). Neuronal synchrony measures. Scholarpedia 2, 1347. doi: 10.4249/scholarpedia.1347

Hennequin, G., Agnes, E. J., and Vogels, T. P. (2017). Inhibitory Plasticity: Balance, Control, and Codependence. Annu. Rev. Neurosci. 40, 557–579. doi: 10.1146/annurev-neuro-072116-031005

Henrie, J. A., and Shapley, R. (2005). LFP Power Spectra in V1 Cortex: The Graded Effect of Stimulus Contrast. J. Neurophysiol. 94, 479–490. doi: 10.1152/jn.00919.2004

Huang, C., Zeldenrust, F., and Celikel, T. (2022). Cortical Representation of Touch in Silico. Neuroinformatics 20, 1013–1039. doi: 10.1007/s12021-022-09576-5

Jensen, O., and Mazaheri, A. (2010). Shaping Functional Architecture by Oscillatory Alpha Activity: Gating by Inhibition. Front. Hum. Neurosci. 4. Available at: https://www.frontiersin.org/articles/10.3389/fnhum.2010.00186 (Accessed April 6, 2023).

Jia, X., Xing, D., and Kohn, A. (2013). No Consistent Relationship between Gamma Power and Peak Frequency in Macaque Primary Visual Cortex. J. Neurosci. 33, 17–25. doi: 10.1523/JNEUROSCI.1687-12.2013

Jiang, H.-J., Qi, G., Duarte, R., Feldmeyer, D., and Albada, S. J. van (2023). A Layered Microcircuit Model of Somatosensory Cortex with Three Interneuron Types and Cell-Type-Specific Short-Term Plasticity. 2023.10.26.563698. doi: 10.1101/2023.10.26.563698

Kondo, S., and Ohki, K. (2016). Laminar differences in the orientation selectivity of geniculate afferents in mouse primary visual cortex. Nat. Neurosci. 19, 316–319. doi: 10.1038/nn.4215

Lagzi, F., and Fairhall, A. L. (2024). Emergence of co-tuning in inhibitory neurons as a network phenomenon mediated by randomness, correlations, and homeostatic plasticity. Sci. Adv. 10, eadi4350. doi: 10.1126/sciadv.adi4350

Lee, J. H., Whittington, M. A., and Kopell, N. J. (2013). Top-Down Beta Rhythms Support Selective Attention via Interlaminar Interaction: A Model. PLOS Comput. Biol. 9, e1003164. doi: 10.1371/journal.pcbi.1003164

Litwin-Kumar, A., Rosenbaum, R., and Doiron, B. (2016). Inhibitory stabilization and visual coding in cortical circuits with multiple interneuron subtypes. J. Neurophysiol. 115, 1399–1409. doi: 10.1152/jn.00732.2015

Luz, Y., and Shamir, M. (2016). Oscillations via Spike-Timing Dependent Plasticity in a Feed-Forward Model. PLOS Comput. Biol. 12, e1004878. doi: 10.1371/journal.pcbi.1004878

Mejias, J. F., Hernandez-Gomez, B., and Torres, J. J. (2012). Short-term synaptic facilitation improves information retrieval in noisy neural networks. Europhys. Lett. 97, 48008. doi: 10.1209/0295-5075/97/48008

Mejias, J. F., and Longtin, A. (2012). Optimal Heterogeneity for Coding in Spiking Neural Networks. Phys. Rev. Lett. 108, 228102. doi: 10.1103/PhysRevLett.108.228102

Mejias, J. F., and Longtin, A. (2014). Differential effects of excitatory and inhibitory heterogeneity on the gain and asynchronous state of sparse cortical networks. Front. Comput. Neurosci. 8. Available at: https://www.frontiersin.org/articles/10.3389/fncom.2014.00107 (Accessed January 16, 2023).

Mejias, J. F., Murray, J. D., Kennedy, H., and Wang, X.-J. (2016). Feedforward and feedback frequency-dependent interactions in a large-scale laminar network of the primate cortex. Sci. Adv. 2, e1601335. doi: 10.1126/sciadv.1601335

Mejias, J. F., and Torres, J. J. (2009). Maximum Memory Capacity on Neural Networks with Short-Term Synaptic Depression and Facilitation. Neural Comput. 21, 851–871. doi: 10.1162/neco.2008.02-08-719

Mejias, J. F., and Wang, X.-J. (2022). Mechanisms of distributed working memory in a large-scale network of macaque neocortex. eLife 11, e72136. doi: 10.7554/eLife.72136

Miller, E. K., Lundqvist, M., and Bastos, A. M. (2018). Working Memory 2.0. Neuron 100, 463–475. doi: 10.1016/j.neuron.2018.09.023

Moreni, G., Pennartz, C. M. A., and Mejias, J. F. (2024). Cell-type-specific firing patterns in a V1 cortical column model depend on feedforward and feedback-driven states. 2024.04.02.587673. doi: 10.1101/2024.04.02.587673

Olsen, S. R., Bortone, D. S., Adesnik, H., and Scanziani, M. (2012). Gain control by layer six in cortical circuits of vision. Nature 483, 47–52. doi: 10.1038/nature10835

Onorato, I., Tzanou, A., Schneider, M., Uran, C., Broggini, A., and Vinck, M. (2023). Distinct roles of PV and Sst interneurons in visually-induced gamma oscillations. 2023.04.08.535291. doi: 10.1101/2023.04.08.535291

Papadopoulos, L., Battaglia, D., and Bassett, D. S. (2022). Controlling collective dynamical states of mesoscale brain networks with local perturbations. doi: 10.48550/arXiv.2208.12231

Perrenoud, Q., Pennartz, C. M. A., and Gentet, L. J. (2016). Membrane Potential Dynamics of Spontaneous and Visually Evoked Gamma Activity in V1 of Awake Mice. PLOS Biol. 14, e1002383. doi: 10.1371/journal.pbio.1002383

Potjans, T. C., and Diesmann, M. (2014). The Cell-Type Specific Cortical Microcircuit: Relating Structure and Activity in a Full-Scale Spiking Network Model. Cereb. Cortex 24, 785–806. doi: 10.1093/cercor/bhs358

Ray, S., and Maunsell, J. H. R. (2010). Differences in Gamma Frequencies across Visual Cortex Restrict Their Possible Use in Computation. Neuron 67, 885–896. doi: 10.1016/j.neuron.2010.08.004

Renart, A., de la Rocha, J., Bartho, P., Hollender, L., Parga, N., Reyes, A., et al. (2010). The Asynchronous State in Cortical Circuits. Science 327, 587–590. doi: 10.1126/science.1179850

Rudy, B., Fishell, G., Lee, S., and Hjerling-Leffler, J. (2011). Three groups of interneurons account for nearly 100% of neocortical GABAergic neurons. Dev. Neurobiol. 71, 45–61. doi: 10.1002/dneu.20853

Schneider, M., Broggini, A. C., Dann, B., Tzanou, A., Uran, C., Sheshadri, S., et al. (2021). A mechanism for inter-areal coherence through communication based on connectivity and oscillatory power. Neuron 109, 4050–4067.e12. doi: 10.1016/j.neuron.2021.09.037

Sjöström, P. J., Turrigiano, G. G., and Nelson, S. B. (2001). Rate, Timing, and Cooperativity Jointly Determine Cortical Synaptic Plasticity. Neuron 32, 1149–1164. doi: 10.1016/S0896-6273(01)00542-6

Spaak, E., Bonnefond, M., Maier, A., Leopold, D. A., and Jensen, O. (2012). Layer-Specific Entrainment of Gamma-Band Neural Activity by the Alpha Rhythm in Monkey Visual Cortex. Curr. Biol. 22, 2313–2318. doi: 10.1016/j.cub.2012.10.020

Thomson, A. M., West, D. C., Wang, Y., and Bannister, A. P. (2002). Synaptic Connections and Small Circuits Involving Excitatory and Inhibitory Neurons in Layers 2–5 of Adult Rat and Cat Neocortex: Triple Intracellular Recordings and Biocytin Labelling In Vitro. Cereb. Cortex 12, 936–953. doi: 10.1093/cercor/12.9.936

Tiesinga, P., and Sejnowski, T. J. (2009). Cortical Enlightenment: Are Attentional Gamma Oscillations Driven by ING or PING? Neuron 63, 727–732. doi: 10.1016/j.neuron.2009.09.009

Tremblay, R., Lee, S., and Rudy, B. (2016). GABAergic Interneurons in the Neocortex: From Cellular Properties to Circuits. Neuron 91, 260–292. doi: 10.1016/j.neuron.2016.06.033

Tsodyks, M., Pawelzik, K., and Markram, H. (1998). Neural Networks with Dynamic Synapses. Neural Comput. 10, 821–835. doi: 10.1162/089976698300017502

Tsodyks, M. V., and Markram, H. (1997). The neural code between neocortical pyramidal neurons depends on neurotransmitter release probability. Proc. Natl. Acad. Sci. 94, 719–723. doi: 10.1073/pnas.94.2.719

Turrigiano, G. G. (2008). The Self-Tuning Neuron: Synaptic Scaling of Excitatory Synapses. Cell 135, 422–435. doi: 10.1016/j.cell.2008.10.008

van Kerkoerle, T., Self, M. W., Dagnino, B., Gariel-Mathis, M.-A., Poort, J., van der Togt, C., et al. (2014). Alpha and gamma oscillations characterize feedback and feedforward processing in monkey visual cortex. Proc. Natl. Acad. Sci. 111, 14332–14341. doi: 10.1073/pnas.1402773111

Veit, J., Hakim, R., Jadi, M. P., Sejnowski, T. J., and Adesnik, H. (2017). Cortical gamma band synchronization through somatostatin interneurons. Nat. Neurosci. 20, 951–959. doi: 10.1038/nn.4562

Vinck, M., Bos, J. J., Van Mourik-Donga, L. A., Oplaat, K. T., Klein, G. A., Jackson, J. C., et al. (2016). Cell-Type and State-Dependent Synchronization among Rodent Somatosensory, Visual, Perirhinal Cortex, and Hippocampus CA1. Front. Syst. Neurosci. 9. Available at: https://www.frontiersin.org/articles/10.3389/fnsys.2015.00187 (Accessed July 11, 2023).

Vinck, M., and Bosman, C. A. (2016). More Gamma More Predictions: Gamma-Synchronization as a Key Mechanism for Efficient Integration of Classical Receptive Field Inputs with Surround Predictions. Front. Syst. Neurosci. 10. Available at: https://www.frontiersin.org/articles/10.3389/fnsys.2016.00035 (Accessed July 11, 2023).

Vogels, T. P., Sprekeler, H., Zenke, F., Clopath, C., and Gerstner, W. (2011). Inhibitory Plasticity Balances Excitation and Inhibition in Sensory Pathways and Memory Networks. Science 334, 1569–1573. doi: 10.1126/science.1211095

Wang, X.-J. (1999). Synaptic Basis of Cortical Persistent Activity: the Importance of NMDA Receptors to Working Memory. J. Neurosci. 19, 9587–9603. doi: 10.1523/JNEUROSCI.19-21-09587.1999

Watt, A. J., and Desai, N. S. (2010). Homeostatic plasticity and STDP: keeping a neuron’s cool in a fluctuating world. Front. Synaptic Neurosci. 2. doi: 10.3389/fnsyn.2010.00005

Wilson, H. R., and Cowan, J. D. (1972). Excitatory and Inhibitory Interactions in Localized Populations of Model Neurons. Biophys. J. 12, 1–24. doi: 10.1016/S0006-3495(72)86068-5

Wood, K. C., Blackwell, J. M., and Geffen, M. N. (2017). Cortical inhibitory interneurons control sensory processing. Curr. Opin. Neurobiol. 46, 200–207. doi: 10.1016/j.conb.2017.08.018

Zou, L., Moreni, G., Pennartz, C. M. A., and Mejias, J. F. (2024). Efficient laminar-distributed interactions and orientation selectivity in the mouse V1 cortical column. 2024.11.05.621826. doi: 10.1101/2024.11.05.621826

